# Ablation of telomerase reverse transcriptase in *Leishmania major* results in a senescent-like phenotype and loss of infectivity

**DOI:** 10.1101/2023.11.10.566596

**Authors:** Mark Ewusi Shiburah, Beatriz Cristina Dias de Oliveira, Habtye Bisetegn, Débora Andrade Silva, Luiz Henrique de Castro Assis, Rubem Menna Barreto, Marcos Meuser Batista, Maria de Nazaré Correia Soeiro, Benedito D. Menozzi, Helio Langoni, Juliana Ide Aoki, Adriano Capellazzo Coelho, Maria Isabel N. Cano

## Abstract

The lack of efficient human vaccines and effective nontoxic drugs for leishmaniasis necessitates a search for new therapeutic targets. The telomere environment could provide potential targets against leishmaniasis. TERT, the telomerase reverse transcriptase component, has been on the radar for new therapeutic options against several diseases for more than two decades. In this study, we constructed a full deletion (*Lm*TERT-/-) and an ORF disruption (*Lm*N420) of the gene encoding the TERT component of *Leishmania major*. *Lm*TERT-/- and *Lm*N420 parasites showed replicative and proliferative defects, growth impairment, cell cycle alterations, increased DNA damage, and progressive telomere shortening. Blockage of parasite altruism and the presence of autophagosomes characteristic of a senescent-like phenotype were also detected. *Lm*TERT-/- and *Lm*N420 parasites caused either micro lesion development or no visible lesions in mouse footpads and reduced infectivity in macrophages. While our checks to see if telomere erosion had reached the *SCG* genes involved in lipophosphoglycan modification showed no changes, our proteomic assessment revealed a downregulation of a metacyclic-associated protein. Complementation of the knockout lineages using the WT *Lm*TERT restored some of the lost phenotypes. Therefore, we speculate that the pleiotropic effects of the loss of *Lm*TERT advance the case for using it as a drug target against the parasite.

## Introduction

Over a million new cases of leishmaniasis are reported annually across 98 territories and countries [1–4]. New therapeutics are crucial due to the absence of licensed vaccines and the inefficiency of currently available disease control and treatment methods [5–8]. The search for an alternative therapeutic strategy in treating leishmaniasis is thus a matter of global public health concern [1,5,9,10].

Telomeres are essential to most eukaryotic chromosomes, including *Leishmania* spp., and are crucial to genome stability and survival [11–16]. A proper understanding of the mechanisms and roles of telomeres in parasite survival would provide answers in the search for new and parasite-specific drug targets [17–20]. *Leishmania* spp. telomeres comprise the conserved tandem ‘TTAGGG’ sequence elongated by telomerase [21–23]. The telomerase component of *Leishmania* is mainly composed of telomerase RNA (TER) and telomerase reverse transcriptase (TERT) [24–29]. *Leishmania* TERT conserves all the canonical structural and functional domains found in most TERT but presents amino acid substitutions specific to the genus [24,26,30–33]. An active TERT adds new telomeric repeats to the chromosome ends. This activity of TERT helps to prevent the incidence of genomic instability, apoptosis, and cell senescence, among many other cell defects due to TERT inactivity [27,34–38].

In this study, we set out to inactivate the TERT in *Leishmania major,* a causative agent of tegumentary leishmaniasis, and assess the impact of this manipulation on the parasite’s biology. We generated two distinct TERT mutants, one bearing the full knockout of both *TERT* alleles (denoted *Lm*TERT-/-) and the other, an endogenous deletion of portions of the open reading frame (ORF) of the *TERT* alleles (denoted *Lm*N420). We generated both mutants using the CRISPR-Cas9 approach outlined by Beneke and colleagues [39]. The results presented here undoubtedly show that TERT ablation significantly influenced several biological processes in the parasite, including cell proliferation, telomere shortening, blockage of parasite altruism, and autophagy, leading to a senescence-like phenotype and loss of *in vivo* and *in vitro* infectivity capacity [40,41]. We assessed whether progressive telomere shortening would affect the telomeric copies of *SCG* genes involved in lipophosphoglycan modifications [42,43]. However, our preliminary results showed that most of the *SCG* genes were unaffected. Interestingly, we found differential expression in proteins, including downregulation of a META domain protein. Together, our results advance the case for using *Lm*TERT as a therapeutic target.

## Results

### Deletion of whole and parts of *LmTERT* is deleterious to Leishmania major

Given the role of TERT in other eukaryotes and the recommended checklist for essential genes provided by Wang and colleagues [44], we hypothesized that deleting the entire *TERT* gene may lead to the immediate death of the parasite. Hence, the strategy was to attempt a complete and partial deletion of the *L. major TERT* gene using the CRISPR‒Cas9 knockout system (S1 and S2 Figs) [39,45,46]. We used the *Lm*007 lineage (S1A-S1C Figs) to generate both knockout lineages (S2A Fig). In both deletion strategies, we selected different clones and expanded them into clonal-population lineages. We phenotypically studied the different clonal lineages and chose one of each to continue the experiments since they showed similar growth and telomere length profiles. Successful *Lm*TERT deletion was confirmed using PCR, Southern blotting, and Sanger sequencing (S2B-S2C Figs).

We checked for phenotypic changes and found that the depletion of *Lm*TERT induced growth defects in the parasite. Compared to the control (*Lm*007), *Lm*TERT-/- and *Lm*N420 promastigotes at passages 7 and 20 showed lower cell density in the logarithmic and early stationary growth phase. This growth defect was particularly evident in the delayed time for the knockout parasites to reach the stationary growth phase (Fig 1A). The growth defect was permanent even after 20 passages. Between passages 7 and 20, the parasites showed a consistent and significant difference in growth on days 2, 3, and 4 (Fig 1A). We further evaluated the proliferative capacity of the parasites by incubating log-phase parasite cultures with EdU (5-ethynyl-2’-deoxyuridine). After EdU labeling, most knockout lineages were identified as not committed to replication (Fig 1B, right panel). Under 50% of *Lm*TERT-/- and *Lm*N420 promastigotes at passages 7 and 20 were proliferating after EdU incorporation, against approximately 80% of proliferating cells in the control, *Lm*007 (Fig 1B, left panel).

Considering the importance of the surveillance system in cell growth [20,47–49], it was necessary to see how much of an influence the ablation of *Lm*TERT has on the cell-cycle progression of the parasites, which may also explain the observed growth and replication impairments. We processed promastigote forms with propidium iodide. Data obtained on the DNA content at the log and stationary phases (Fig 1C) revealed changes in the cell cycle progression pattern of knockout lines compared to the control. Knockout cells (*Lm*N420 and *Lm*TERT-/-) showed measurable delays or transient arrest in the G0/G1 phase at the log phase (Fig 1C). At both P7 and P20, a significant number of knockout parasites, *Lm*N420 (50.88% and 58.05%) and *Lm*TERT-/-(48.06% and 52.26%), respectively, were recorded at the log G0/G1 phase versus *Lm*007 (44.87% and 53.91%) (Fig 1C, Panel 1). In contrast, lower percentages of cells were recorded for the knockout lineages at the log G2/M phase, *Lm*N420 (20.50% and 20.22%) and *Lm*TERT-/-(23.18% and 17.75%) against *Lm*007 (27.03% and 20.22%) (Fig 1C, Panel 2). These data suggest an interruption in the cell cycle progression of the knockout parasites and a possible defective transitioning system that could impact parasite development [48].

In the absence of TERT, we expected an increase in DNA damage. This theory was founded on the observed telomere attrition (see Fig 4) and the detected delays in cell cycle progression (Fig 1C). At the log phase of growth, the parasites generally had low DNA damage signals averaging 4.5% for *Lm*007, 0.2% for *Lm*N420, and 3.5% for *Lm*TERT-/-. At the stationary growth phase, this increased drastically to about 38.16% and 36.24% for the TERT-depleted parasites (*Lm*N420 and *Lm*TERT-/-, respectively) compared to an average of 22.48% for *Lm*007 (Fig 1D). Here, we establish an association between the observed DNA damage and the recorded delays in cell cycle progression in the knockout lineages.

**Fig 1.**
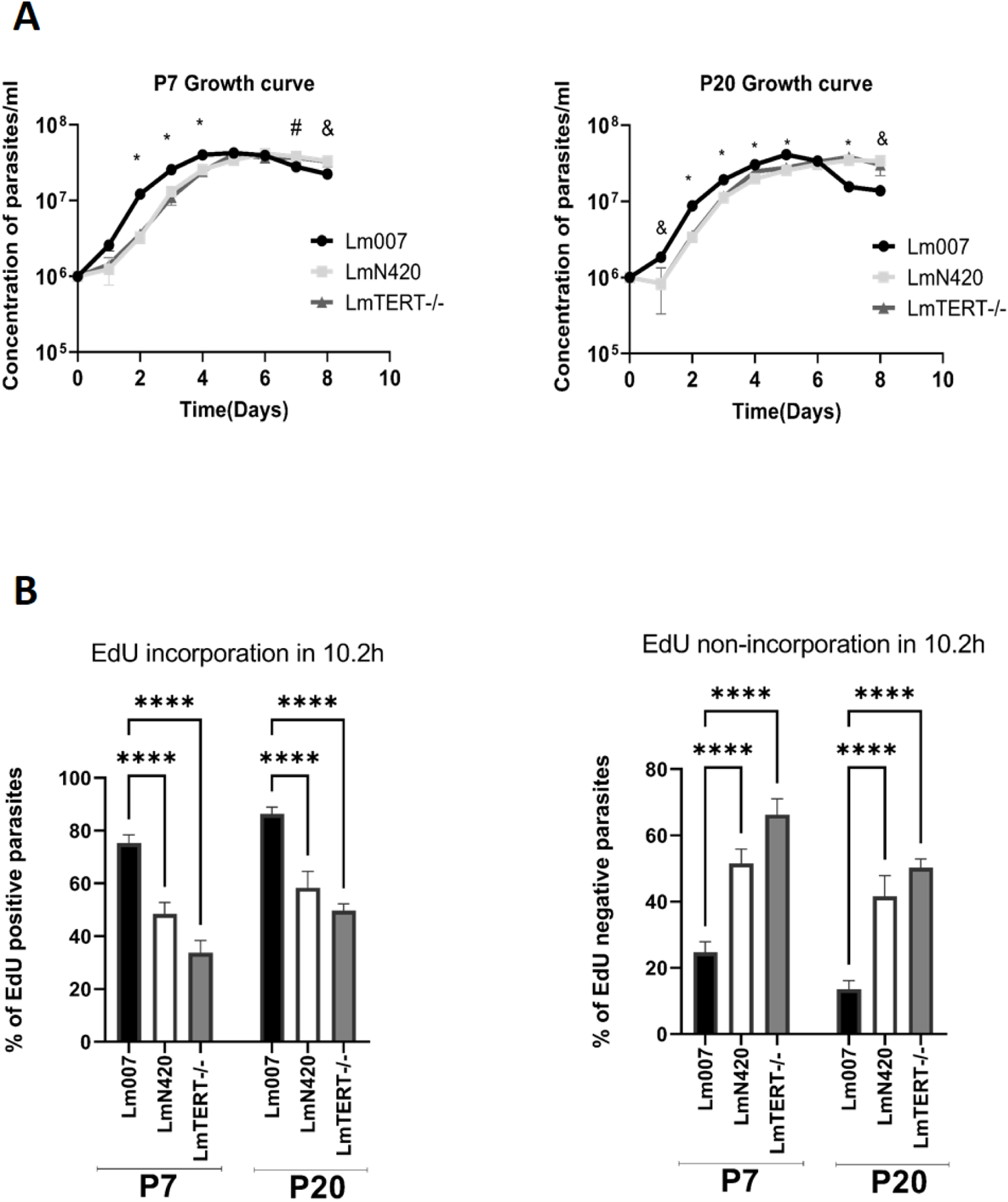
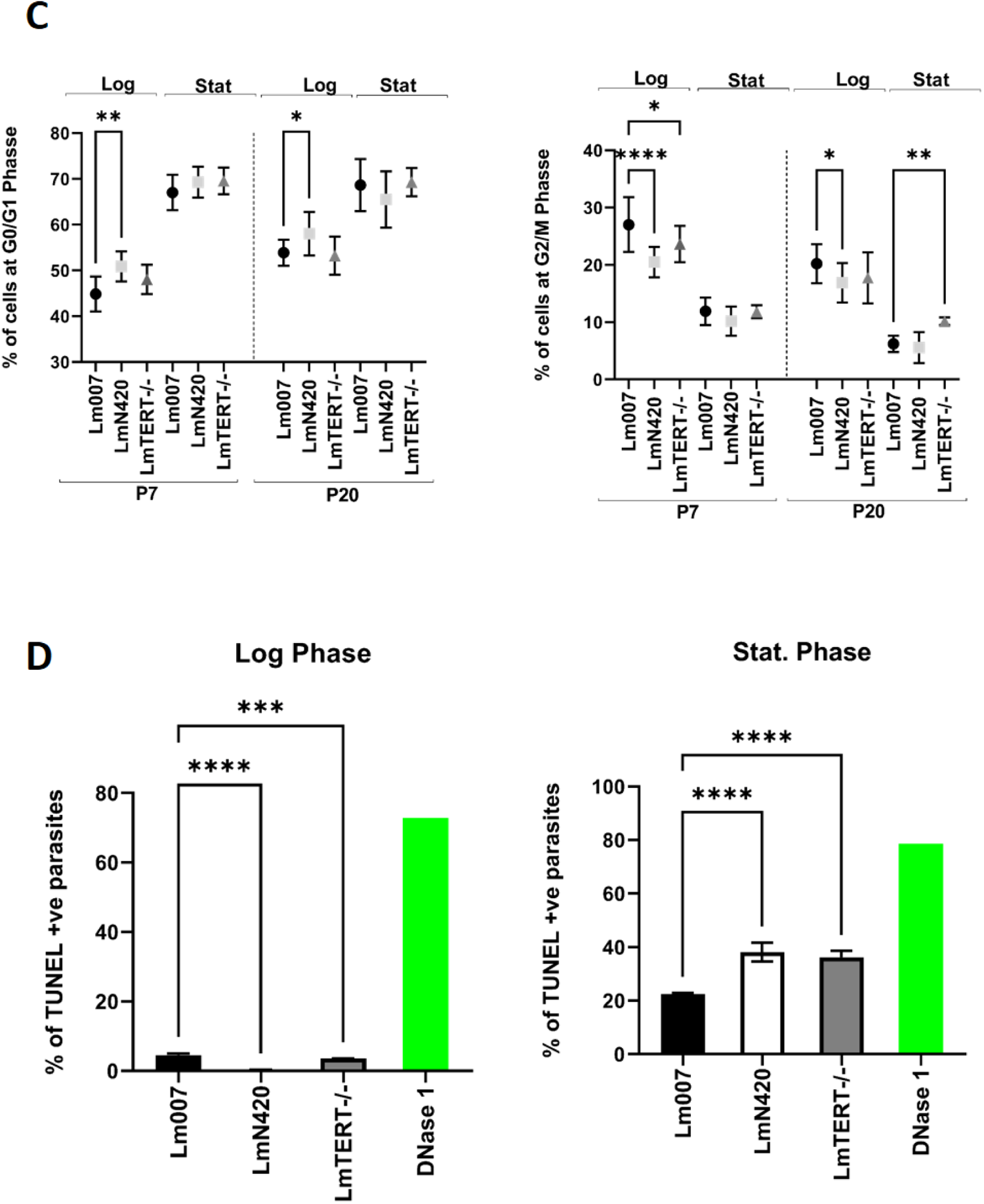
Defective growth, DNA synthesis, and accumulation of DNA damage in TERT-depleted parasites. (A) Growth curves comparing the growth profile of TERT-depleted parasites and *Lm*007 at P7 and P20. The following symbols represent the significant differences between treatments: *, Both edited lines differ from *Lm*007; &, only *Lm*N420 is significantly different from *Lm*007; #, only *Lm*TERT-/-is significantly different from *Lm*007. Statistical analysis was performed using the Welch t-test on the means of three parallel experiments for each treatment group. (B) Parasites’ ability to synthesize DNA was represented by their EdU uptake (left panel) and nonresponsive parasites to DNA synthesis (right panel). Ordinary one-way ANOVA was used as the statistical treatment for five technical replicates from two independent experiments for each group. (C) Graphs from data on parasite challenges in cell cycle progression (D) TUNEL data showing accumulation of DNA damage in parasites after TERT depletion. Data from a single experiment with five technical replicates was used in the ordinary one-way ANOVA statistical test. All data in this Fig is represented as mean ±SD, and significance was tested at P≤0.05.

### TERT-deficient parasites lacked cell death and had an increased number of stationary phase promastigotes negative for peanut lectin agglutinin (PNA)

One of the clauses given by the Nomenclature Committee on Cell Death (NCCD) in declaring cell death is the compromise in the plasma membrane, which symbolizes a point of no return for such cells [50,51]. We used an Annexin-PI assay to check parasite death by membrane turnover and permeabilization. The control parasites (*Lm*007) showed lower viability records (56.78% and 62%) at P7 and P20, respectively, when compared with *Lm*N420 (94.40% and 95.16%) and *Lm*TERT-/-(91.50% and 94.14%) (Fig 2A). In addition, *Lm*007, in contrast to the TERT-depleted parasites (*Lm*N420 and *Lm*TERT-/-), was nearly tenfold more PI/Annexin-positive, showing an average of 37.72% versus 3.83% and 6.64% in *Lm*N420 and *Lm*TERT-/-, respectively, at P7 (Fig 2B).

We performed a metacyclic selection assay to ascertain the parasites’ normal development. We collected parasites grown for up to 7 days at the stationary phase for P7 and P20 and performed PNA agglutination [52]. The results presented in Fig 2C show a surprisingly high number of promastigotes negative for PNA in the knockout lineages. Statistical analysis revealed a significant difference for *Lm*N420 and *Lm*TERT-/-against *Lm*007, as shown in the left and right panels of Fig 2C, respectively. Additionally, the *Lm*TERT-/-remarkably quadrupled its previous count at P20 and was fourfold higher than *Lm*007 at P20 (Fig 2C, right panel). At the stationary phase, *Leishmania* promastigotes cultures should have more Annexin-V-positive cells and fewer parasites negative for PNA [53,54]. The unexpectedly low permeability, Annexin-V binding of the knockout parasites at the stationary phase, and the increased number of PNA-negative parasites suggest a modification in the parasites’ plasma membranes after TERT ablation.

**Fig 2.**
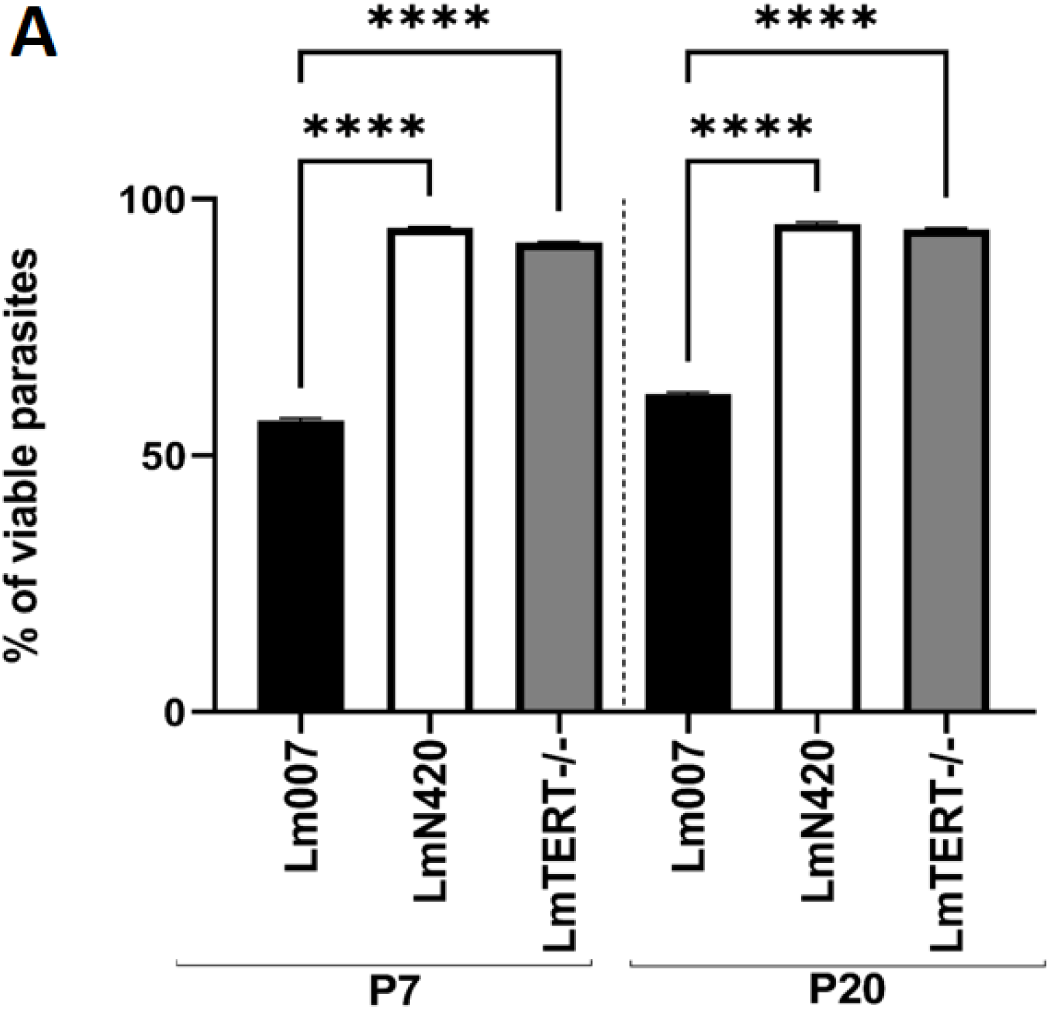
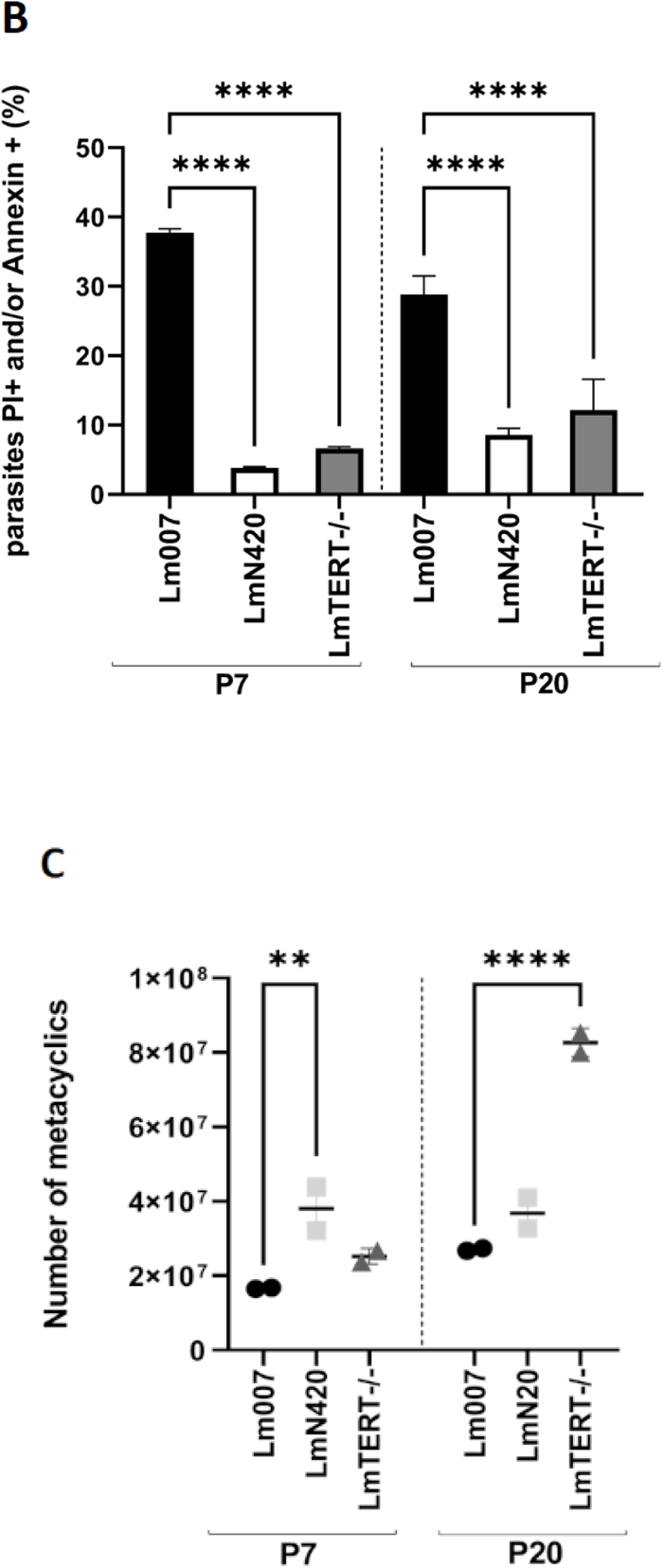
Absence of cell death and increased PNA-negative parasites among the TERT-depleted *L. major* promastigotes. (A) and (B) Parasites at the stationary phase were stained with Annexin-V and propidium iodide to determine viability. (A) The graph represents parasites that are positive for both stains, indicative of no cell death (quadrant-Q4, Fig. S3C). An FMO control was performed by treating *Leishmania* parasites with only PBS, and no staining procedure was conducted on these parasites to represent an intact membrane. (B) The graph shows the percentages of parasites found in quadrants Q2 and Q3, Fig. S3C. Positive control was performed by treating *Leishmania* parasites with 20% formaldehyde for approximately 10 min to cause membrane turnover before being subjected to a staining procedure. The negative control was parasites with no stain. (C) The graph shows the number of parasites that are negative for lectin agglutination. Ordinary one-way ANOVA was used in the statistical treatment of graphs presented in this Fig to establish significance at P≤0.05. Graphs (A) and (B) represent two independent experiments of at least three technical replicates. Graph (C) is representative of two experiments conducted in parallel.

### Ultrastructural analysis of *L. major* wild-type promastigotes and TERT-deficient parasites shows senescent-like phenotypes

To check eventual ultrastructural alterations in TERT-depleted promastigotes, wild-type and mutant parasites of different passages *in vitro* (P5 and P20) were processed for scanning and transmission electron microscopy. Our findings demonstrated a lack of morphological alteration when *L. major* promastigotes (with and without TERT deletion) were examined by scanning electron microscopy (SEM). Parasites presented characteristic morphology in all experimental conditions, such as elongated shapes and free flagella. At the same time, round and intermediate forms could also be observed, in addition to dividing promastigotes (Fig 3, SEM panel), showing a heterogeneous population of cells in culture.

Transmission electron microscopy (TEM) evaluated wild-type promastigotes displaying typical organelles, such as the nucleus, mitochondrion, kinetoplast, endoplasmic reticulum, and flagellum (Figs 3A and 3D). In contrast, TERT-deficient promastigotes at P5 (Figs. 3B and 3E, TEM panels) and P20 (Figs. 3C and 3F, TEM panels) presented cytosolic vacuolization (V) and the formation of autophagosomes and blebs in the plasma, flagellar and flagellar pocket membranes. Also, mitochondrial insults such as organelle swelling and inner concentric membrane structures characteristic of autophagy were noted (Figs. 3B, 3C, 3E, and 3F, TEM panels). The features observed here, the prolonged delays in cell cycles, and the replication troubles detailed earlier are reminiscent of a cell undergoing senescence [40,41,55,56].

**Fig 3.**
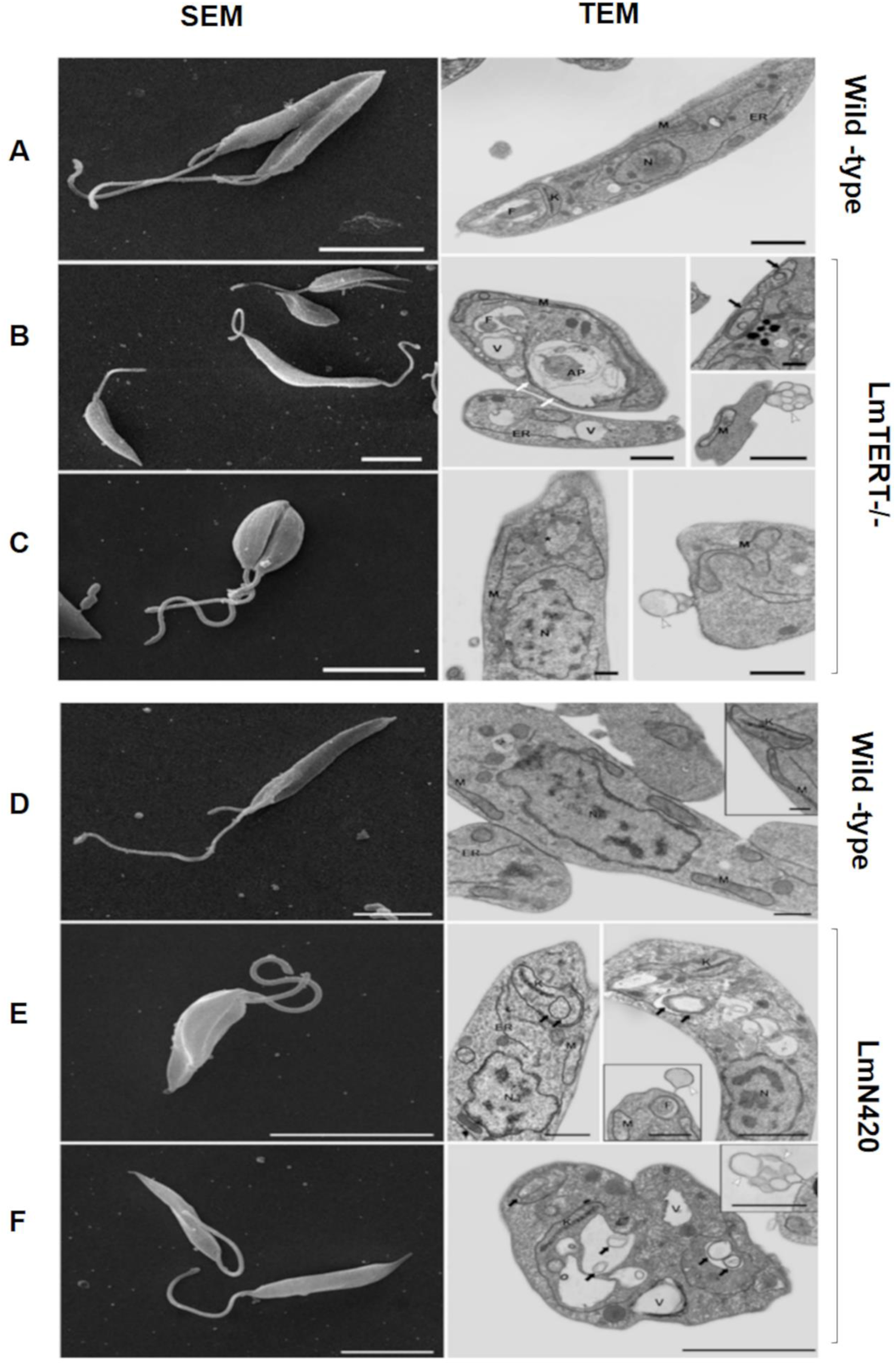
TERT-depleted parasites show a senescent-like phenotype. Left panels (A-F), scanning electron microscopy (SEM) (bars = 5 µm) and Right panels (A-FI), transmission electron microscopy (TEM) (bars = 1 µm) of wild-type, LmTERT-/-, LmN420 promastigotes. In the top and bottom panels, the wild-type parasites (A, D) show typical, elongated morphology and normal appearance of organelles, including the nucleus (N), mitochondrion (M), kinetoplast (K), endoplasmic reticulum (ER), and flagellum (F). In the top panel, promastigotes TERT-/-at P5 (B, SEM) and P20 (C, SEM) show classical elongated bodies and flagella. Their analyses by TEM show cytosolic vacuolization (V), formation of autophagosomes (AP), blebs in the flagellum and flagellar pocket (white arrowheads), mitochondrial phenotypes such as organelle swelling (*), and the presence of inner concentric membrane structures (black arrows). In the bottom panel, promastigotes LmN420 showed no morphological alteration by SEM obtained at P5 (C) and P20 (F), with parasites presenting characteristic morphology such as elongated shape and free flagellum. Additionally, round and intermediate forms were observed, in addition to dividing promastigotes. When evaluated by TEM, LmN420 promastigotes at P5 (E, TEM) and P20 (F, TEM) presented cytosolic vacuolization, blebs in the plasma, flagellar and flagellar pocket membranes (white arrowheads), and the presence of concentric membrane structures inside mitochondria (black arrows).

### Telomere attrition in TERT-depleted *L. major* promastigotes and recovery after complementation reveals the role of TERT in parasite telomere maintenance

A characteristic feature of most organisms with interrupted TERT activity is a reduction in telomere length [34,41,57]. TERT is responsible for the polymerization of telomeric ends. Its ablation leads to an incomplete telomerase holoenzyme and terminates telomere length maintenance, resulting in continuous attrition of these ends [18,25,58–62]. Whether the TERT in *Leishmania major* has similar roles as in other eukaryotes was evaluated here by Southern telomere restriction fragment (TRF) assessment and Flow-FISH. Telomere attrition was evident in the TERT-depleted parasites at passages 7 and 20 (Figs. 4A and 4B). Proportional changes in telomere length, calculated using arbitrary fluorescence units (a.u.) of molecules of equivalent soluble fluorochrome (MESF) (Table 1), showed an approximately 50% reduction in telomere size after TERT inactivation.

We performed a complementation strategy by introducing an episomal plasmid carrying wild-type TERT back into the respective knockout lineages. We denoted the complementation lineages as NAB and TAB for *Lm*N420 and *Lm*TERT-/-, respectively. The telomere profiles from these lineages revealed a significant increase in telomere size compared to their respective knockouts (Fig 4B, left panel). Approximately 25-30% of the lost telomeres were recovered (Fig 4B, right panel). These data point to the critical role of TERT in maintaining chromosomal termini in these parasites.

**Fig 4.**
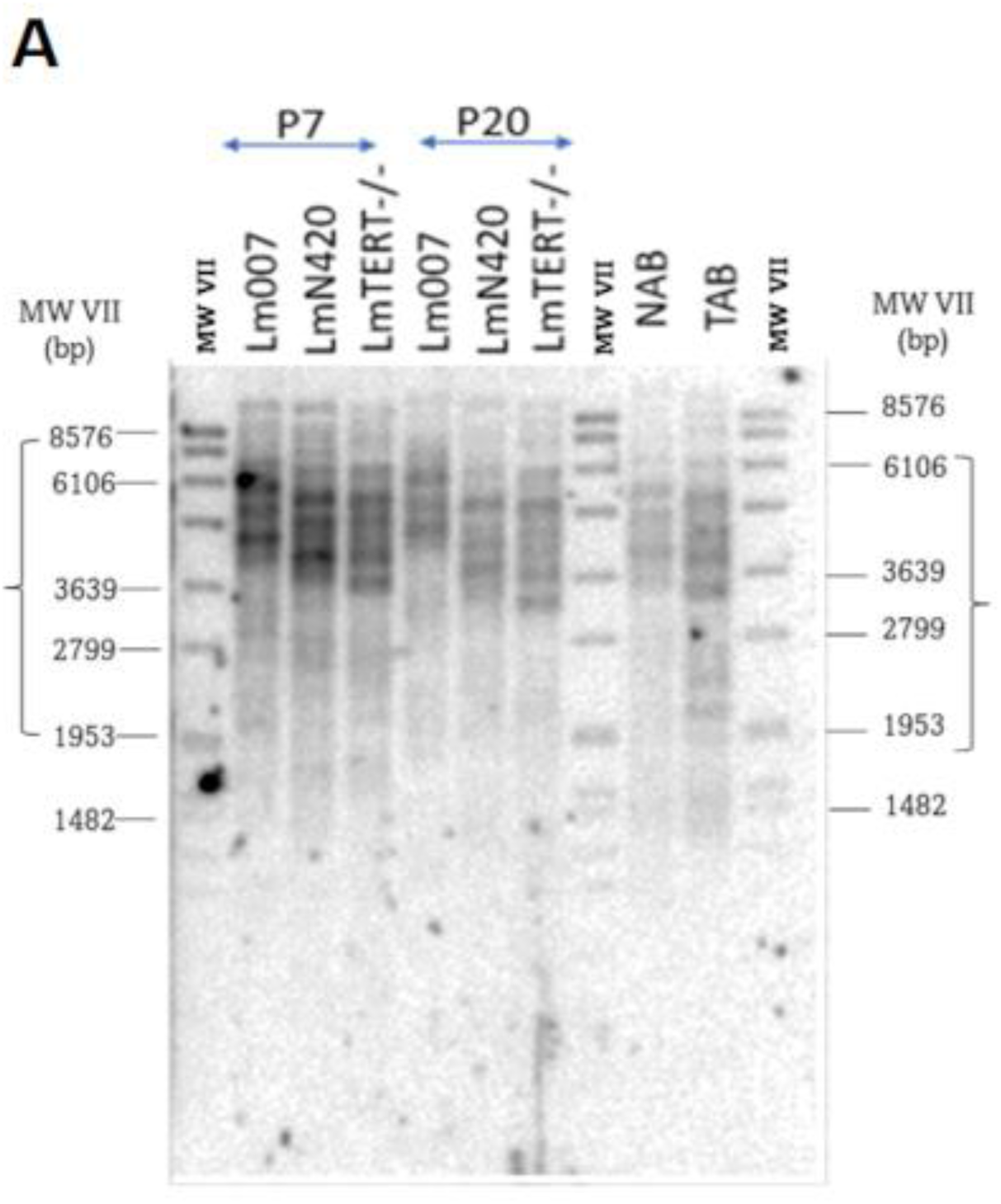
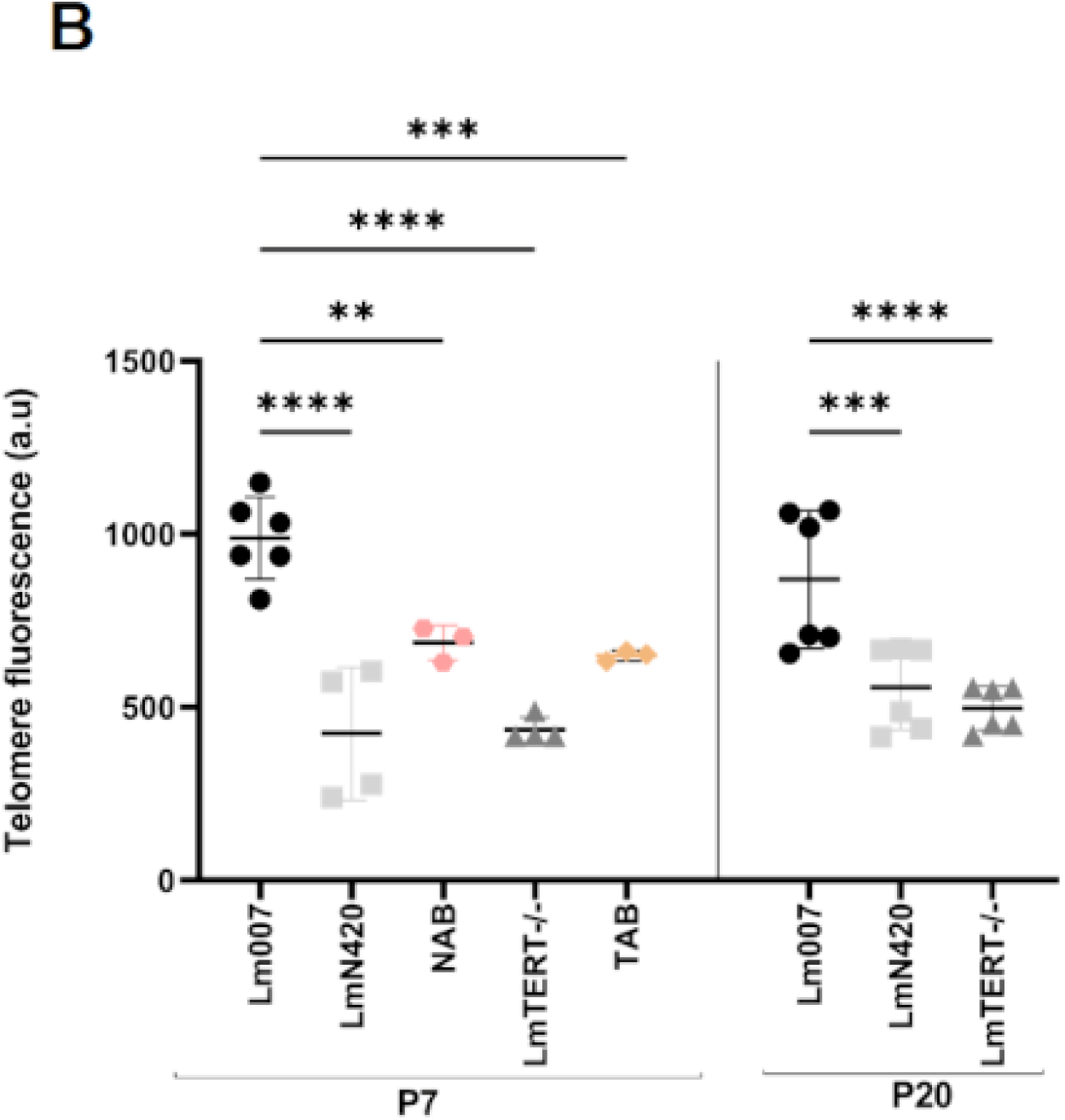
TERT depletion induces telomere attrition in *L. major* promastigotes. The deletion of *Lm*TERT leads to a significant reduction in telomeres. (A) The Southern blot image shows the different groups at passage 7 (left panel), passage 20 (middle panel), and after complementation with the episomal expression of wild-type TERT (NAB and TAB) in the right panel. (B) The fluorescent values were obtained by calculating the MESF for *Lm*007, *Lm*N420, and *Lm*TERT-/-, as well as NAB and TAB at P7 (left panel) and P20 (right panel). Significance was obtained after ANOVA treatment of data obtained from at least three technical replicates of two independent experiments. Graphs are shown with mean ± SD data. P≤0.05.

**Table 1.**
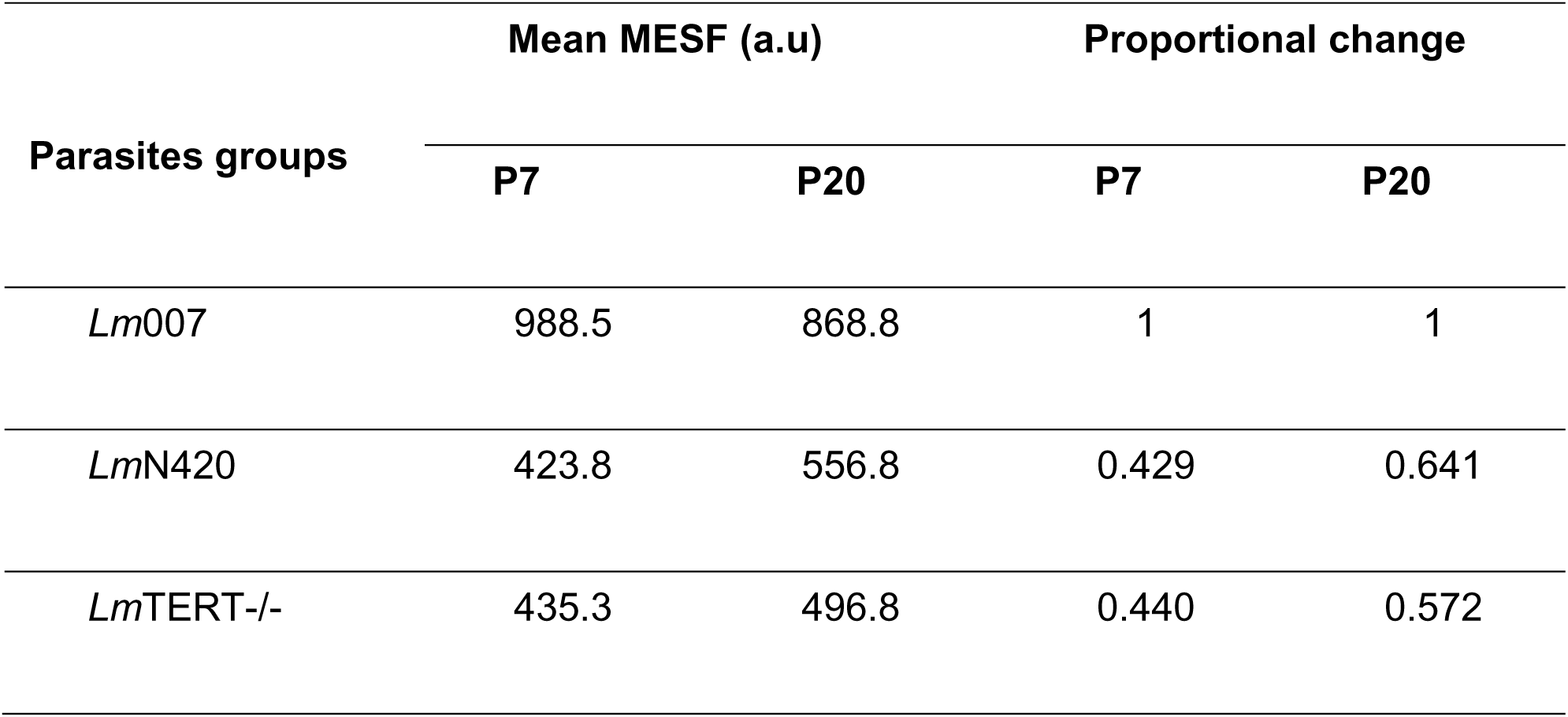
Comparative changes in telomere size.

### TERT-depleted parasites showed reduced or no capacity to infect BALB/c mice models, and bone marrow-derived macrophages from BALB/c

Due to the low Annexin-positive and high number of PNA-negative promastigotes, we questioned the infective capacity of the knockout lineages. We conducted concurrent Annexin-V and *in vivo* infectivity assays (Figs 5A and 5B). An *in vitro* infectivity assay using bone marrow-derived macrophages (BMDMs) from BALB/c mice was also performed at 24 h, 48 h, and 72 h (Figs 5C and 5D). Data from the *in vivo* infection assay showed a significant difference in lesion development between the mice infected with the knockout lineages and the *Lm*007-infected group. All mice in the *Lm*007 treatment group developed their first visible lesion at 50 days post-inoculation. In contrast, mice infected with *Lm*TERT-/-cells had only two out of the three showing microlesions 61 days post-infection. None of the three mice infected with the *Lm*N420 cells presented any lesion development during data collection (61 days post-inoculation). In addition, one mouse from the *Lm*N420 group was experimentally separated for further monitoring. This mouse showed no visible lesions up to 70 days post-inoculation.

The sizes of the inoculated and non-inoculated footpads were measured using Vernier calipers, and the difference was estimated as the swelling size (Fig 5B, graph). At a mean size of approximately 5 mm, the *Lm*007-infected mouse paw was about twice the size observed in *Lm*TERT-/-and that of *Lm*N420. A mean difference of 1.9 mm and 2.5 mm was observed between *Lm*007 versus *Lm*TERT-/-and *Lm*N420, respectively (Fig 5B, graph). Concurrently, serological tests using indirect immunofluorescence (IIFA) reported a negative presence of infection in mice inoculated with the knockout lineages using different titrations. We only obtained positive signals from the experiment’s positive control and the mouse group infected with *Lm*007 (Supplementary Figs 4B and 4C).

In agreement, macrophage infection studies using BMDMs at a parasite-to-macrophage ratio of 10:1 revealed reduced infection in TERT-deficient lineages. At 24 h, the *Lm*007-infected macrophages had less infection compared with the knockout (*Lm*N420 and *Lm*TERT-/-) and complementation lineages (NAB and TAB) (Fig 5C). However, at 48 h, the percentage of macrophages infected with *Lm*007 increased significantly from a mean of 8.357% to 56%. In contrast, the percentage of cells infected with *Lm*N420 declined from 23.47% at 24 h to 13.33% at 48 h. Similarly, infection with *Lm*TERT-/-declined from 27.24% to 7.409%. Complementation lineages, NAB and TAB, progressively increased from 32.18% and 17.33%, respectively, to 57.90% and 19% at 48 h. A continuous increase in the percentage of infected macrophages was recorded for TAB (39%) at 72 h. In comparison, a continuous decline was recorded for the knockout lineages (*Lm*N420 and *Lm*TERT-/-). The decline was observed in the controls *Lm*007 (27.33%) and NAB (56%) at 72 h, which were nevertheless significantly higher than the knockout lineages. An infectivity index, calculated by finding the product of the mean number of amastigotes per macrophage and the percentage of macrophages infected, also revealed a similar pattern of infection as the percentage of macrophages infected (Fig 5D). These results imply that *Lm*TERT influences the infectivity of *Leishmania major* parasites and their growth inside the host cells.

**Fig 5.**
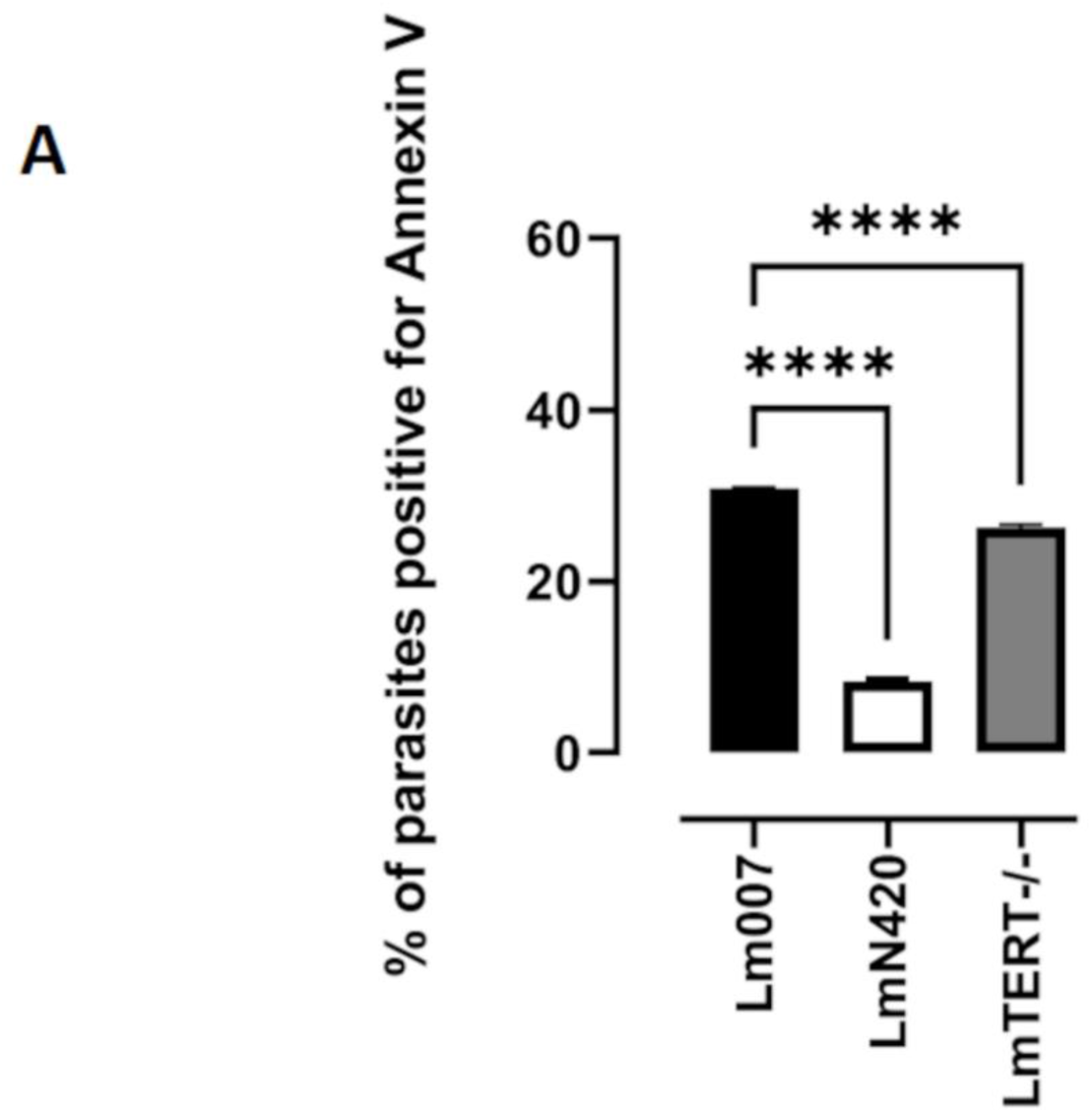
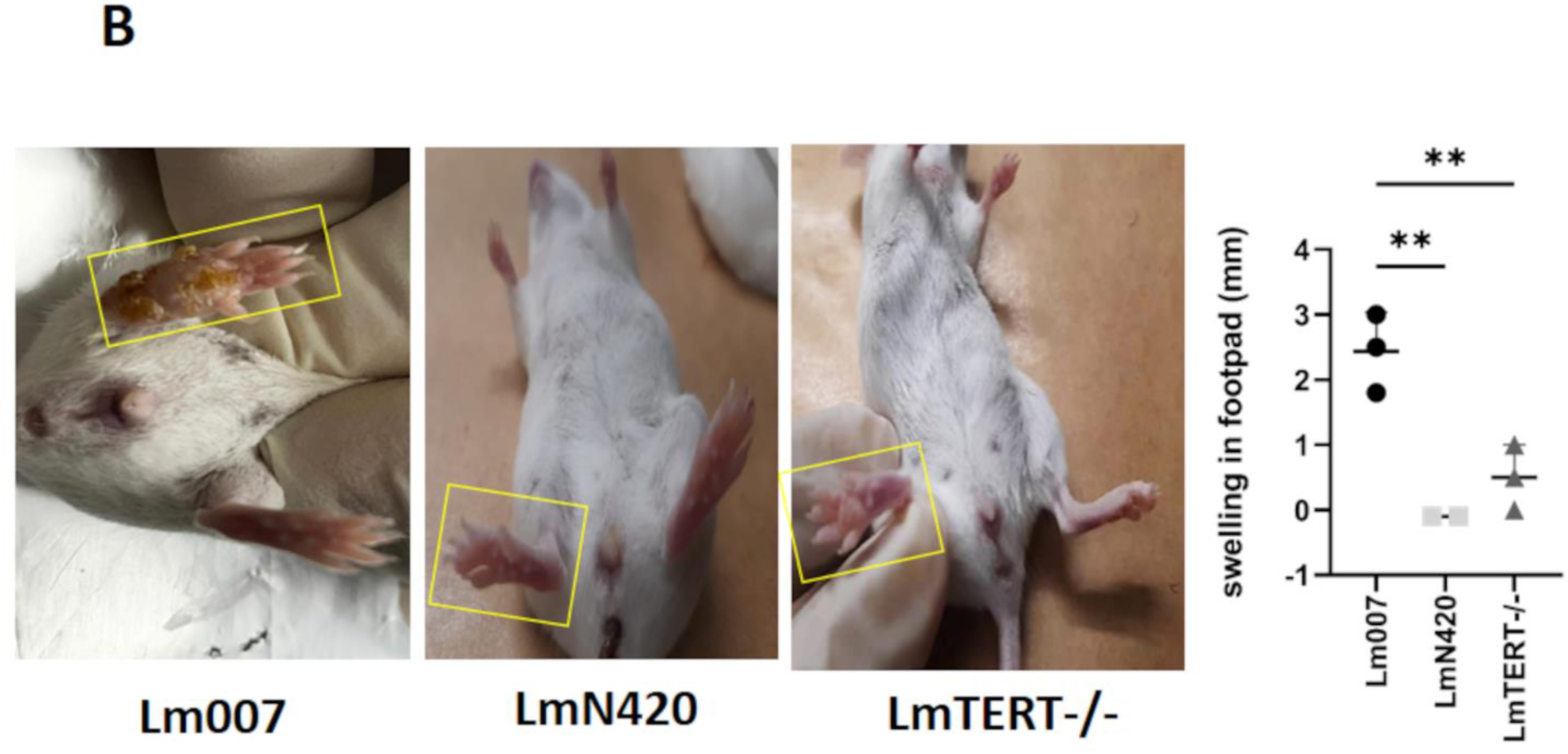
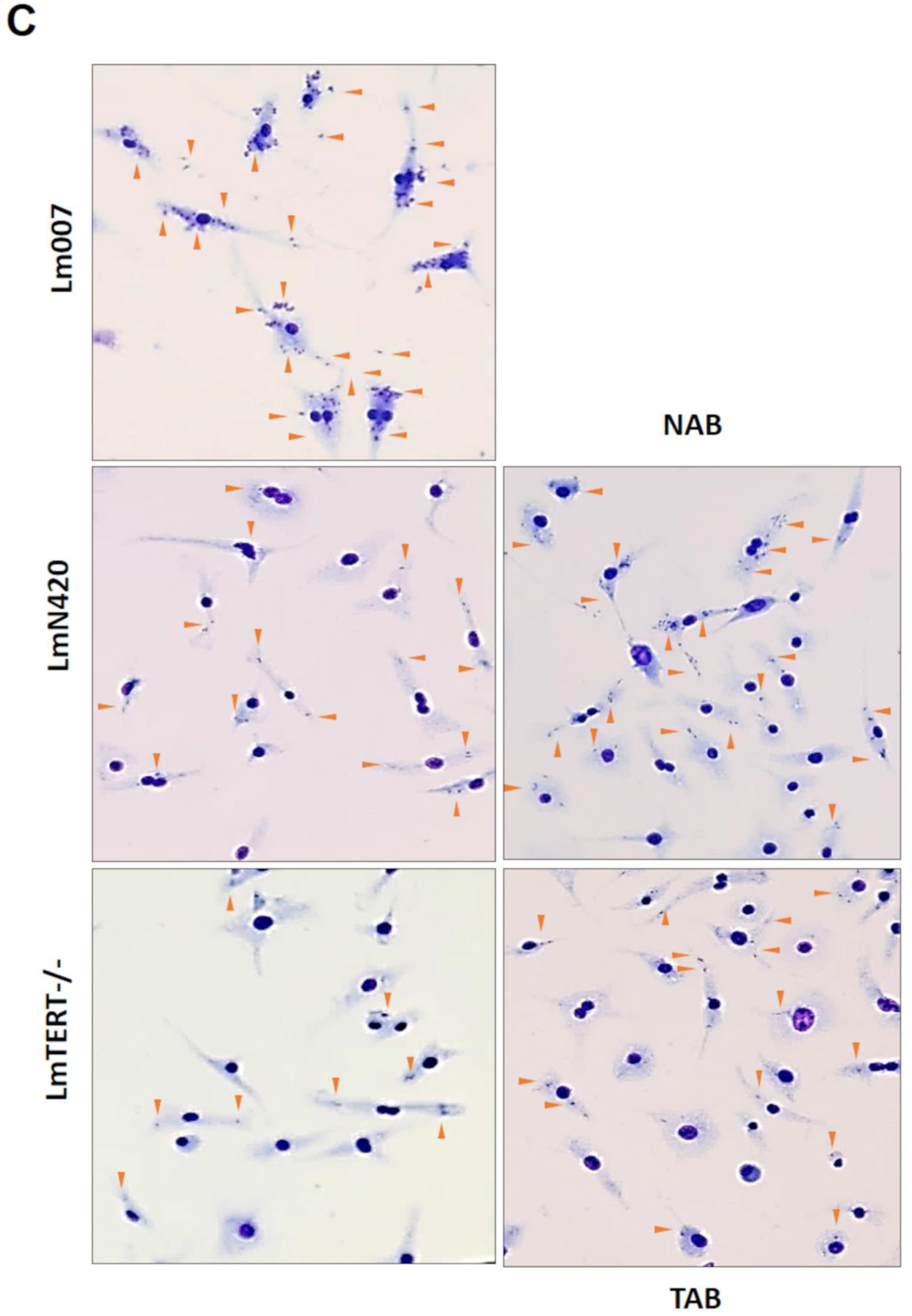
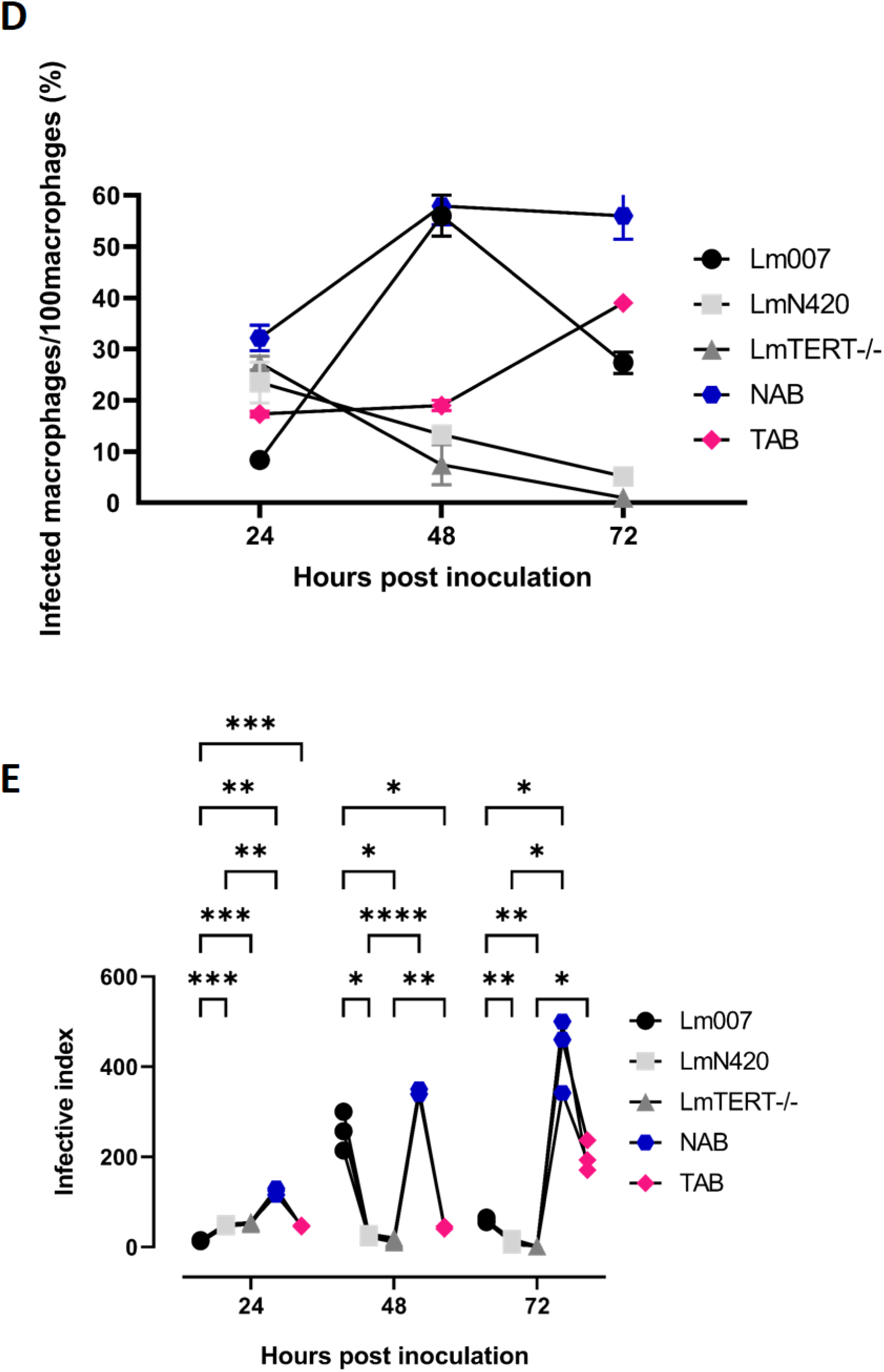
TERT-depleted parasites showed reduced or no capacity to infect BALB/c mouse models and bone marrow macrophages. (A) The graph shows that Annexin assays were conducted concurrently during inoculation using the parasites from the same cultures used in mouse inoculation. Positive control was performed by treating *Leishmania* parasites with 20% formaldehyde for approximately 10 min to cause membrane turnover before taking them through a staining procedure. (B) Lesion size determined after 61 days post-inoculation is represented in the graph. Representative images of mouse footpads infected with *Lm*007, *Lm*N420, and *Lm*TERT-/-are shown from left to right. (C) Images showing some macrophages invaded by amastigotes. Arrows point to the number of amastigotes per macrophage. (D) The graph of macrophage infections performed, including data from complementation parasite infection. (E) This graph shows the infection index and representative images of macrophage infection. Data from approximately 300 macrophages per treatment group (three replicates in parallel) were subjected to two-way ANOVA.

### Telomeric *SCG* genes remain undisturbed

Linking the results observed in the telomere attrition, metacyclic selection assay, and the infectivity of parasites, we checked for the telomeric *SCG* genes involved in lipophosphoglycan (LPG) modification in the parasites. We reasoned that the shortening of telomeres may have impacted these genes, resulting in modifications in their roles as LPG modulators [42,43,63]. Due to the high sequence identity (>90%) between the *SCG* genes, we only designed primers for the sets with less similarity. Performing ordinary PCR with specific primers, we confirmed that telomeric erosion had not reached the point of eroding some of the genes (S5 Fig).

### Mass spectrometry confirms protein deregulation in TERT-depleted parasites

While the *SCG*s amplified were still present, our further analysis of the *Lm*TERT-/-and *Lm*007 proteomes by mass spectrometry showed changes in protein regulation. Differential protein expression analysis revealed 104 proteins (Data file Table 2, S7 Fig) being significantly upregulated or downregulated. Among the downregulated proteins were the stage-regulated metacyclic proteins (META domain-containing protein) (S7B Fig). META domain proteins are expressed less in the knockout compared to the *Lm*007. The proteomic data suggest that the increased PNA-negative promastigote numbers from the metacyclic selection assay (Fig 2C) may not be true metacyclics. Instead, they may be promastigotes whose plasma membrane composition has been altered, thereby affecting the lipophosphoglycan constitution and subsequent agglutination by PNA. Another interesting result in the proteomics assessment was the upregulation of a TPR-repeat (tetratricopeptide repeat) containing protein (S7CFig). TPR motifs are known facilitators of protein-protein interactions. They are present in proteins involved in different biological functions, such as in the orthologues of EST1p (ever-short telomere 1 protein), an evolutionarily conserved regulator of telomerase activity [64,65].

## Discussion

Telomerase reverse transcriptase (TERT) is an essential component of telomerase. Nonetheless, the extent of its relevance to *Leishmania* parasite biology and survival has been unexplored. Consequently, this study sought to understand the effects of TERT deletion on parasite morphology, cell cycle progression, cell proliferation, cell fate, cell organelle organization, infectivity, and telomere homeostasis. We show that the absence of TERT does not lead to the immediate death of parasites as hypothesized but does affect vital parasite biology, including *in vivo* and *in vitro* infectivity and the regulation of parasite surface lipophosphoglycan constitution.

The absence of the TERT in *Leishmania major,* unlike *Trypanosoma brucei,*^60^ shows detrimental effects on parasite growth, causing the cells to reach the stationary phase later than *Lm*007. The delay resulted in the difference in timing where the decline phase for the control parasites happened earlier than the knockouts, likely due to nutrient depletion and other conditions associated with stationary phase cultures, such as changes in the culture media pH [66]. The ability of cells to go through the various stages of development hinges on their transitions across the distinct phases of the cell cycle. In the presence of genomic perturbations, cells activate checkpoints to ensure that the integrity of their genome is uncompromised. In addition, cell cycle checkpoints guarantee that the next generation of cells has the necessary cellular supplies to survive, grow, and reproduce [20,48,67]. Our data indicate that *Lm*TERT-/-and *Lm*N420 are challenging in synthesizing DNA. The data also suggests the likely initiation of checkpoints due to DNA damage leading to arrest in cell cycle progression. From our data, we posit a role of TERT in DNA synthesis and cell cycle progression. Interestingly, these challenges do not seem to hinder the continuous growth of the parasite. Damasceno and colleagues [68] demonstrated that *L. major* could continue with other cell cycle processes even with a checkpoint activation. Our data agree with their observation [68].

The telomere attrition recorded here due to TERT manipulation and a similar report by Dreesen *et al*. [58] on *T. brucei* indicates the significance of TERT in maintaining telomeres in these ancient eukaryotes. The reduction in telomere length signifies the loss of the protective cap at the end of the chromosome, triggering a senescent-like state [19,69]. There are reports of telomerase-deficient cells using alternative telomere maintenance mechanisms for survival [70–73]. These post-senescence survivors activate alternative lengthening of telomere (ALT) strategies, such as recombination and chromosome circularization, to maintain short telomeres [74]. Although not much evidence is provided here, based on the observed phenotypes and concurrent events in other cell types, we suspect that there is a population of our TERT-deficient parasites that use an ALT mechanism to continue to proliferate despite having short telomeres. Telomere uncapping led to an accumulation of DNA damage signals as TERT-deficient parasites grew from the log into the stationary phase [40,54,75–81]. Similar observations have been reported in *Saccharomyces cerevisiae,* whereby telomere uncapping led to the accumulation of DNA damage as cell division proceeded [82].

While the use of Annexin-V as a marker for apoptosis in *Leishmania* has been called into question, some relevant studies have shown a strong relationship between this assay and parasite infectivity and cell fitness. Annexin-V labeling demonstrates changes to the parasite plasma membrane [51,53,83–89]. Based on this, we declare cells as either dead or alive without stressing the type of death [90]. The low levels of Annexin-V-positive cells recorded for *Lm*N420 and *Lm*TERT-/-indicate intact membranes and, thus, viable parasites. The presence of Annexin-V-positive cells in culture has been reported to serve as an altruist, helping in manipulating host cell defenses to facilitate the intracellular survival of the fittest parasite [91]. We suspected abolished infectivity in TERT-ablated parasites based on the Annexin data. This suspicion was confirmed by the *in vivo* and *in vitro* infectivity results. The absence of infection in mice inoculated with *Lm*TERT-/-revealed by the indirect immunofluorescence assay suggests that the microlesion observed in the footpad likely did not develop from infection with the knockout *Leishmania* parasites. While indirect immunofluorescence detection of *Leishmania* is a qualitative measure with the disadvantage of false-negative or positive results, it is an effective method for detecting cutaneous leishmaniasis [92,93]. Furthermore, the controls used in our study show high robustness in the sensitivity and specificity of the tests (S4 Fig). The infectivity index recorded after infecting BMDMs corroborates these observations and incontrovertibly indicates that TERT is involved in parasite infectivity.

In addition, the formation of autophagosomes revealed by electron microscopy results points to the initiation of cellular control by autophagy [90]. Autophagy is a molecular pathway described in *Leishmania* as a cell control mechanism directly involved in survival and infectivity [50,68,90]. Basmaciyan and colleagues [94] have shown that autophagic *Leishmania* parasites have lower Annexin-positive signals, a phenomenon corroborated by our results here. Therefore, it is likely that in the absence of TERT, *Leishmania* parasites use an autophagic mechanism similar to macroautophagy in mammalian cells. Macroautophagy allows cells to break down their cytosolic materials and organelles to produce energy for survival under adverse conditions [94,95]. Given this information, we propose a suggestive role of TERT in parasite pro-survival-autophagy.

Another question is whether the selected metacyclic parasites are actual metacyclics or metacyclic-like’ parasites. Metacyclics are considered a key determinant of *Leishmania* virulence, but the data obtained here are contrary [96]. Despite this observation, Zandbergen and colleagues [54] reported a positive correlation between virulence and Annexin-positive *Leishmania* parasites. The authors consider the presence of Annexin-positive parasites as a better determinant of infection than the number of metacyclic parasites. Our Annexin, infectivity, and METAs results corroborate their report [54]. As a result of the modifications in the plasma membrane and the protein deregulation observed, we believe the promastigotes obtained from TERT-depleted lineages are parasites lacking the necessary potential for PNA binding. *Leishmania* parasites are known to have specific lipophosphoglycan (LPG) at each growth phase. Although our results from the amplification of *SCG* genes do not conclusively suggest that telomeric erosion has affected them, it is impossible to confirm whether their protein expression or function was compromised. Additionally, it is unclear to what extent the other unamplified members of the *SCG* gene family influence the LPGs [42,43].

In conclusion, our study has revealed evidence of *TERT* involvement in maintaining and regulating *L. major* telomeres, parasite infectivity, the cell cycle, surface protein expression, metacyclogenesis, DNA damage machinery, and prosurvival autophagy. We provide several lines of evidence on how the absence of *Lm*TERT leads to pleiotropic effects that can be likened to senescence in yeast and other cells. We propose a model in which the absence of *Lm*TERT leads to telomere shortening, thereby triggering cell cycle arrest and DNA damage responses that cause cells to enter a replicative senescent state. We suspect these cells survive using prosurvival autophagic mechanisms and maintain short telomeres using an unknown ALT mechanism. Future studies will try to elucidate the exact ALT mechanism utilized by these parasites. On the other hand, the TPR-containing protein overexpressed by the TERT-/-parasites can be a starting point by determining its relatedness and relevance in comparison with that of the EST1-TPR in *S. cerevisiae,* which was found to be relevant in telomere homeostasis [65]. The consistent results from the full-gene knockout and the ORF-disrupted knockout, along with the complementation results, further strengthen the notion that the absence of TERT was responsible for the observed pleiotropic defects. As such, *Lm*TERT can serve as a drug target against leishmaniasis.

## Materials and methods

### Resource availability statements Materials availability

*Leishmania* mutant lines and oligonucleotides generated in this project are available upon request to the lead contact.

### Data availability

All data reported in this paper can be found at doi:10.5281/zenodo.10035576, along with the respective statistical treatments and results obtained.

### Experimental model and subject details

WT *Leishmania major* Friedlin strain (MHOM/IL/1980/FRIEDLIN) was used to generate the *Lm*007 lineage, which was used to generate knockout lineages. Knockout lineages served as background lines for the respective complementation lineages. Experiments were conducted at two key passages, passage 7 and passage 20. Depending on the assay, these experiments used logarithmically growing parasites and stationary phase parasites. All parasites were cultivated with the M199 (Cultilab) culture medium supplemented with 10% Foetal Bovine Serum, 41.75 mM Hepes (Amresco), 0.35 g/l NaHCO_3_ (Sigma), 104.38 µm Adenine (Sigma), 0.001% Biotin (Sigma), 10000 U/mL penicillin/streptomycin solution and kept at 26 °C [97]. For the selection of single, independent clones, promastigotes were plated on solid M199 media in the presence of 20% FBS mixed 1:1 with 2% Agar (bacteriological grade) supplemented with specific antibiotics. The resistance gene carried by the parasite lineage determined the choice of antibiotics. Female Balb/c mice at six weeks old were also used in the experiment. Mice were fed adlibitum and housed in an authorized facility.

### Gene editing

#### Design and synthesis of oligonucleotides

SgRNAs were obtained following similar strategies described by Beneke *et al.* [45] and Yagoubat *et al.* [98]. A set of oligonucleotide sequences for the gene ID *Lmj*F.36.3930 (*Lm*TERT component) was designed using information from the <u>LeishGEdit</u> (http://www.leishgedit.net/mysql/primersearch.php). Another oligonucleotide set was designed using the Eukaryotic Pathogen CRISPR guide RNA/DNA Design Tool (EuPaGDT) (http://grna.ctegd.uga.edu/), maintaining default parameters. Whereas LeishGEdit sgRNA targeted the UTR regions of the gene, the EuPaGDT-designed sgRNAs targeted the endogenous region of the gene, 420bp and 4022bp from the start codon for the 5’ and 3’ cut, respectively. Two sets of oligonucleotides were designed from the <u>LeishGEdit site</u> to amplify a DNA fragment from a pTplasmid [39]. Two fragments of plasmids that confer resistance to different selection drugs (neomycin and puromycin) were used to amplify a donor template (DT). Puromycin and Neomycin donor templates were used in the full TERT knockout replacement, whereas only the Neomycin donor template was used in the *Lm*N420 deletion.

#### Parasite transfections and clonal lineage selection

The Cytomix buffer, composed of 120 mM KCl, 0.15 mM CaCl_2_, 10 mM K_2_HPO_4_, 25 mM HEPES, 2 mM EDTA, and 2 mM MgCl_2_, was used in the deletion procedures. Briefly, 2×10⁸ cells were centrifuged at 1000 g for 10 min, washed with the buffer at a similar speed, and the pellet resuspended in 500 µL of the buffer. To generate *Lm*N420, NAB, *Lm*007, and TAB, two pulses from the Bio-Rad gene pulser system were used with the following conditions: 25 µF and 1500 Volts (3.75 kV), with 10 s wait period between the pulses [97]. The Amaxa nucleofector system (Lonza) generated the *Lm*TERT-/-; a single pulse of the X-001 program was used [45,68]. Clonal lineages were selected by plating in the presence of specific antibiotics. Using two different systems to generate knockout lineages increased the confidence in attributing observed phenotypes to the gene manipulation rather than a possible confounding effect of the system.

### Flow cytometry techniques

#### Assessment of DNA content

A total of 5 × 10^6^ cells were harvested from log phase and stationary phase cultures followed by resuspension in 90% methanol, incubated at -20 °C for 30 min. Two PBS washes were performed on permeabilized cells before staining with 40 μg/mL propidium iodide and 100 μg/mL RNAse A (Amresco). Stained cells were incubated at 37 °C for 30 min, then at 4 °C for at least 30 min before collection with a flow cytometer (BD Biosciences C6 Acuri). 20-30,000 events were collected in specific PE-A and PE-H gates for further deconvolution or data analysis using FlowJo vX 10.0.7r2 (Tree Star, Inc., Ashland, OR). During analysis, events between treatments were maintained within a difference of 1,000 events. The same gating strategy with adjustments to capture an equal number of events in each instance [99–101]

#### EdU labeling of parasites

20 μM of EdU (Sigma) was added to the culture medium at the exponential growth phase [102]. Cells were incubated for 10.2 h in the EdU before collection for processing. Cells were permeabilized and incubated in the click-IT reaction cocktail for 30-45 min, then were washed and resuspended in 200 μl of propidium iodide buffer before, incubated at 37 °C for 30 min and later 4 °C for 30 min. A similar procedure for collection and analysis was followed, as in DNA content analysis above [103–106].

#### Annexin-V and cell membrane integrity assessment

1 x 10^6^ stationary phase parasites were collected, washed with PBS, and resuspended in 100 μl of Binding buffer, 2 μl of Annexin V, and 100 μg/mL of Propidium Iodide (Bio-Rad). The mix was incubated at room temperature for 15 min before analysis. Channels FITC-A and PE-H were used because this Annexin-V is conjugated to FITC 20,000-30,000 single events were collected using the Flow cytometer, and data analysis was performed using FlowJo vX 10.0.7r2 (Tree Star, Inc., Ashland OR) [53].

#### Flow-FISH telomere length assessment

2 x 10^7^ cells were collected and then permeabilized with 90% Methanol at -20 °C overnight. Hybridization buffer was added to controls for non-detection, and for the treatment, 300 μl of Hybridization buffer containing TelC probe (1:10,000) was added. The cells are flickered and incubated at 85 °C for 15 min, followed by overnight room-temperature incubation. Cells were washed and incubated at 40 °C, followed by resuspension in PI solution and incubated at 37 °C for 30 min before being transferred to 4 °C for 30 min. 10,000 to 20,000 events were collected for analysis. MESF values were calculated using the manufacturer-recommended program.

#### Tunel evaluation of DNA damage

2 x 10^7^ parasites were fixed with 4% paraformaldehyde (Sigma) and 0.2% Triton X-100 permeabilization. Permeabilized cells were equilibrated and incubated with the nucleotide mix and Terminal Deoxynucleotidyl Transferase, Recombinant (rTdT) enzyme (Promega). Cells were then incubated at 37 °C for 60 min, protected from direct light, followed by treatment with 0.1% Triton (Bio-Rad) containing 5mg/mL BSA(Promega) dissolved in PBS. Procedures for staining cells for the assessment of DNA content were followed. Finally, 10,000-20,000 events were collected by flow cytometry.

### Transmission (TEM) and scanning electron microscopy (SEM) analysis

*Leishmania major* procyclic promastigotes of *Lm*007, *Lm*N420, and *Lm*TERT-/-each at passages 7 and 20 were washed twice with 0.1 M phosphate buffer saline (PBS) and fixed at 4 °C for 40 min with 2.5% glutaraldehyde (GA) diluted in 0.1 M Na-cacodylate buffer (pH 7.2). For TEM analysis, the parasites were washed three times with 0.1M sodium cacodylate buffer and post-fixed for 15 min with 1% osmium tetroxide (OsO_4_), 0.8% potassium ferricyanide and 5 mM CaCl_2_ in 0.1 M cacodylate buffer, pH 7.2. Then, samples were washed with the same buffer, dehydrated in an ascending acetone series, and embedded in PolyBed 812 resin as previously described [107]. Ultrathin sections were stained with uranyl acetate and lead citrate and examined under a Jeol 1200 EX transmission electron microscope (Tokyo, Japan) at Centro Nacional de Biologia Estrutural e Bioimagem (CENABIO). Alternatively, for SEM analysis, fixed parasites adhered on glass coverslips coated with 0.1% poly-L-lysine and post-fixed as described above. Then, samples were dehydrated in a crescent ethanol series (30-100 %), dried using the critical point method with CO2, mounted on aluminum stubs coated with a 20 nm gold layer, and examined with a Jeol JSM6390LV scanning electron microscope (Tokyo, Japan) Platform Rudolf Barth in Instituto Oswaldo Cruz, FIOCRUZ [108].

### Metacyclic selection

Approximately 1 x 10^8^ cells of stationary phase parasite culture were centrifuged at 2,300 x g for 5 min and washed twice with PBS 1X. The supernatant was discarded, and metacyclic solution (PBS1X, 0.1 mg/mL of lectin PNA (Sigma), and 5 mM EDTA) was added to the samples. The mixture was allowed to incubate at room temperature for 1 h, and the supernatant was collected as metacyclics by low-speed centrifugation [52,96].

### Bacterial transformation and confirmation of *Lm*TERT presence

Thermo-competent XL-1 blue bacteria were incubated with 10 ng of plasmid for 30 min. After the incubation period, the sample was transferred to a prewarmed incubator at 42 °C for 2 min and set on ice for 5 min. SOB media was supplemented with 100 μl of 1 M MgCl_2_, 100 μl of 1 M MgSO_4_, and 900 μl was added to the samples before setting to incubate rotating at 37 °C for an hour. A solid plate of LB agar was prepared, and aliquots of the transformed bacteria were cultured overnight. A single colony from the overnight incubation was selected and expanded in LB broth by overnight incubation at 37 °C. After the overnight expansion, the protocol for Midi-Prep extraction of DNA was followed as directed by the manufacturer to extract the DNA of the plasmid. Restriction fragment analysis was done using *Cla*1 and *Pst*1 restriction enzymes to confirm the extracts.

### DNA and RNA extraction

DNA extraction was performed using the Qiagen kit and following the protocol provided by the manufacturer. About 1 x 10^8^ parasites from passages 7 and 20 were taken through the protocol. The final elute of the DNA was RNase-treated before quantification. Proteinase K and RNase treatment were performed at 56 °C and 37 °C, respectively. For RNA extraction, about 1 x 10^8^ cells were treated with Trizol (Invitrogen) to extract the RNA. RNA and DNA were quantified by BioTek Epoch Microplate Spectrophotometer (BioTek), and quality checks were done by EtBr-stained agarose gel electrophoresis.

### PCR, Sanger sequencing, and RT-qPCR

PCRs were performed using either the Taq polymerase (Invitrogen) in reactions requiring downstream usage of PCR products or the GoTaq green master mix (Promega), where downstream application was not required. A primer walk strategy was designed along the edited regions using SnapGene V.3.2.1. We amplified the edited regions of the genome. PCR products were purified using the PureLink™ Quick Gel Extraction and PCR Purification Combo Kit (Invitrogen). The purified PCR products were Sanger sequenced at a core facility (IBTEC). Assembly of contigs from fragments and further analysis were done using Geneious V.7.1.3.

To perform RT-qPCR, 1µg of extracted RNA from passage 7 was DNase (Thermo scientific) treated and converted to cDNA using the iScript™ cDNA Synthesis Kit (Bio-Rad). Primers for *Lm*TERT (target) and RPN8 (reference) were used to check the expression of the reintroduced, wildtype *Lm*TERT in knockout lineages by qPCR. The primer sequences are listed in the S1 Table. qPCR reactions were performed using a StepOnePlus™ Real-Time PCR System (Applied Biosystems). Each reaction contained a 2.5 ng cDNA template, 100 nM of forward and reverse primers, 5 μl SYBR Green PCR Master Mix (Bio-Rad), and nuclease-free water. The qPCR cycling conditions consisted of an initial denaturation at 95 °C for 10 min, followed by 40 cycles of denaturation at 95 °C for 15 sec, annealing at 60 °C for 30 sec, and extension at 72 °C for 30 sec. The threshold cycle (Ct) values were determined for each sample, and relative gene expression levels were calculated using the 2∧-ΔΔCt method, with normalization to the reference genes (RPN8). All qPCR experiments were performed in triplicate, and the results are presented as mean ±SD.

### Terminal restriction fragments profile obtained by telomeric Southern blot

Genomic DNA extracted from the parasites at passages 7 and 20 were digested with 5 U *Rsa*1 (Roche) at 37 °C overnight. Terminal restriction fragments (TRF) were fractionated in a 0.8% EtBr-stained agarose gel and transferred to the Hybond N+ nylon membrane (Cytivia) overnight. The membrane was crosslinked after the transfer and taken through stringent washes before being hybridized with a 5’-DIG-labelled telomeric probe (TELC) (5’-DIG-CCCTAACCCTAACCCTAACCCTAACCCTAA). The hybridization signals were developed with an anti-DIG-HRP conjugate antibody and CSPD-Star (Roche).

### Protein studies

#### Protein extraction and western blot

Whole-cell lysates from parasites at P7 were obtained by washing twice with phosphate-buffered saline (PBS) at 3500 x g for 5 min. 1 mL of Buffer A (20 mM Tris-HCl, 1 mM EGTA, 1 mM EDTA, 15 mM NaCl) supplemented with 1 mM spermidine, 0.3 mM spermine, and 1 mM DTT was added to cells and incubated in liquid nitrogen for 1 min. Tubes were recovered from liquid nitrogen and placed on ice to defrost. 0.5 % Nonydet p-40 and protease inhibitor were added to the extract and centrifuged at maximum speed for 20 min. The supernatant was recovered as total protein extract, and the pellet was discarded. 10 % SDS-polyacrylamide gel was prepared, and samples were loaded into wells in the gel and subjected to electrophoresis to separate the bands in two steps. Samples were run against a standard molecular weight marker at 80 V, 400 mA, and 400 W for 1 h in the first step, then at 120 V, 400 mA, and 400 W for another hour in the second step. Transfer to Immun-Blot® PVDF membrane for protein blotting (Bio-Rad) was done in a Tris-glycine transfer buffer (25 mM tris and 192 mM Glycine) overnight at 90 mA and 120 V at 4 °C evenly cooled temperature. A blocking solution was prepared using a blotting-grade blocker (Bio-Rad) at a 1% (w/v) concentration with PBS/TWEEN 20 (PBST) solution. Primary antibodies were prepared in the blocking solution at appropriate dilutions. Following the manufacturer’s instructions, the ECL™ western blotting analysis system (GE Healthcare) was used on the membrane to generate signals (Genesys V.1.7.2.0).

#### Proteomic analysis

The protein extracts of promastigote forms of *Lm*TERT-/-and *Lm*007 were obtained following a modified version of the protocol described by Fragaki *et al.* [109], using 1 x 10^9^ promastigotes on the last day of the exponential phase. Briefly, the cells were collected by centrifugation at 2500 x g for 5 min at 4 °C and washed twice with 1 mL of sterile phosphate-buffered saline (PBS) using the same parameters for the centrifugation steps. The washed cells were resuspended in 500 µL of lysis buffer 1, which was composed of 10 mM Hepes, 1.5 mM MgCl2, 10 mM KCl, 0.5 mM dithiothreitol (DTT), 0.5% NP40, and 1x protease inhibitors cocktail (Sigma-Fast-Ethylenediaminetetraacetic acid (EDTA)-free). This first lysis step aimed to release the cytoplasmatic proteins, and after the cell resuspension by vortex, the microtube was kept on ice for 30 min. The cytoplasmatic proteins in the supernatant were recovered by centrifugation at 2000 x g for 10 min at 4 °C. The nuclear proteins were further obtained in two steps. First, the pellet was treated with 50 U of DNAse I (Thermo Scientific) for 20 min at room temperature in 300 mM Hepes pH 7.5, 8 mM MgCl_2_, 200 mM KCl, and 1,5 x protease inhibitors cocktail. Then, 50 µL of lysis buffer 2 (120 mM Hepes pH 7.5, 1.5 M KCl, 500 mM sucrose, 0.4 mM EDTA, and 1 mM DTT) was added, and the microtube was incubated on ice for 20 min. Nuclear proteins were recovered by centrifugation at 16100 x g for 15 min at 4 °C. The protein extracts were quantified using PierceTM B.C.A. Protein Assay kit (Thermo Fischer Scientific) according to the manufacturer’s instructions. Approximately 300 µg of proteins were submitted to a precipitation step using acetone/methanol. After resolubilization, 50 µL of 50 mM Hepes pH 7.9 protein concentrations were obtained using the same B.C.A. procedure mentioned above. For the in-solution protein digestion procedure, 100 µg of total protein was denatured in 7 M urea. The disulfide bonds were reduced with 10 mM DTT for 40 min at room temperature, and cysteine residues were carbamidomethylated by incubation with 40 mM of iodoacetamide for 40 min at room temperature. The excess of iodoacetamide was consumed following incubation with 10 mM DTT for 10 min at room temperature. For proteolysis, the protein solutions were diluted to 700 mM urea with 50 mM Hepes pH 7.5, and aliquots containing 1 µg of proteins were saved as the non-digested sample for posterior comparison. Trypsin (Sigma) was added in a 1:50 (m/m) ratio, and the reactions were incubated at 30 °C for 18 h. After proteolysis, the solutions were acidified with formic acid and concentrated by vacuum centrifugation. Then, the peptides were desalted employing the Stage Tip approach with C18 membranes (3M) and following the procedure described by Rappsilber et al. [110]. The desalted peptides were dried by vacuum centrifugation, dissolved in 30 µL of 0.1% formic acid, and quantified using the Qubit TM Protein Assay kit (Thermo Fischer Scientific). The volume containing 10 µg of peptides was dried by vacuum centrifugation and labeled with TMT as described by Zecha et al. [111]. The reactions were split into cytosolic and nuclear proteins and followed the same labeling scheme: the fractions from *Lm*007 were labeled with tag 127N, and those from *Lm*TERT-/-were labeled with tag 128N.

#### LC-MS/MS analysis

The peptide fractions (cytosolic and nuclear) were dissolved in 0.1% formic acid (FA), and 100 ng was analyzed by LC-MS/MS. Samples were loaded onto a 2 cm precolumn (nanoviper C18, 3 μm) in an EASY-nLC 1200 system (Thermo Fischer Scientific). The liquid chromatography was performed using a 15 cm C18 analytical column (nanoviper C18, 2 μm). The peptide elution followed a gradient of 0.1% FA as phase A and 100% ACN/0.1% FA as phase B with a constant flow rate of 300 nL/min. The starting condition was 5% B, reaching 28% B in 80 min, 10 min to 40% B, and finally, the gradient was increased to 95% B in 2 min, remaining in this condition for 12 min. The nLC system was in tandem with an Orbitrap Fusion Lumos mass spectrometer (Thermo Fischer Scientific) operated in positive mode. The automatic gain control (A.G.C.) was set to 4 x 10^5^ ions and a maximum fill time of 50 ms, while the mass range was fixed to 400-1600 m/z with high resolution (120,000 full-width half maximum (FWHM) at m/z 200). Data-dependent acquisition was selected, and a cycle time of 3 seconds was achieved. Higher energy collision-induced dissociation (H.C.D.) was applied for peptide fragmentation using 38 as normalized collision energy (N.C.E.). The fragmentation step was performed at high resolution (50,000 FWHM) with an A.G.C. target of 1 x 10^5^ and a maximum injection time of 200 ms using an isolation window of 0.7 m/z. The MS/MS selection intensity threshold was set to 2.5 x 10^4^ and a dynamic exclusion of 60 s.

#### Data analysis

Database search was performed using MaxQuant software (version 2.0.3.0) [112] and in collaboration with Professor André Zelanis (UNIFESP). All raw files of LC-MS/MS of the two peptide fractions (cytosolic and nuclear) were submitted to database search against a fasta file of protein sequences from *L. major* (Friedlin 2021) available in TriTrypDB [113] (8508 sequences in March 2023). Database search was carried out considering the 10plex TMT labeling as the type of analysis. The reporter ion analysis was specified at the MS2 level, considering a mass tolerance of 0.003 Da, and the ‘direct mode’ was selected. The precursor ion fraction (P.I.F.) was set to 0.75, and no normalization method was applied. Trypsin was chosen as the enzyme in a specific mode. As dynamic modifications, the occurrence of methionine oxidation (+15.9949 Da) and deamidation of N.Q. (+0.9840), and Protein N-term acetylation (+42.0106) were considered. Cysteine carbamidomethylation was considered a fixed modification (+57.0215). The search mode “match between runs” was enabled in all searches. A false discovery rate (F.D.R.) < 1% was set at P.S.M. (Peptide Spectrum Match) and protein levels. The results were analyzed using the Perseus software (version 2.0.9.0). Only protein groups with two or more peptides were considered as identified.

### Infectivity studies

#### Inoculating mice with *Leishmania major*

Parasites were cultured until they reached the stationary phase (day 7). BALB/c Female mice at 6 weeks old were brought in from a mice facility and nurtured until 7 weeks 4 days when they were inoculated with 40 μl of PBS containing about 2 x 10^6^ of parasites from *Lm*007, *Lm*TERT-/-and *Lm*N420 in suspension. Inoculation was performed in the right hind footpad of each mouse for all treatment groups using a needle and syringe. Feeding and water were provided ad libitum, with regular monitoring and changes in bedding material. Three mice were used per treatment, and lesion development was followed. Mice were euthanized 61 days post-inoculation. 100 μl of blood was collected before being euthanized. Both hind footpads were measured, and the size of the swelling was determined. The study was conducted under the guidelines for the care and use of laboratory animals.

#### Macrophage derivation from bone marrow and infection

Bone marrow-derived macrophages (BMDMs) were obtained from femurs and tibias of BALB/c mice. The cells were incubated in RPMI 1640 medium supplemented with penicillin (100 U/mL), streptomycin (100 µg/mL), L-glutamine (2 mM), sodium pyruvate (1 mM), 20% fetal bovine serum and 20% L929 conditioned medium as macrophage stimulating factor source, for 7 days at 37 °C and 5% CO_2_. After differentiation, BMDMs were collected by centrifugation at 600 x g for 10 min at 4°C and counted in the Neubauer chamber. 3 x 10^5^ cells were incubated in RPMI medium, as described in sterile coverslips (Olen) in 24-well plates (Nest) overnight at 37°C and 5% CO_2_. Non-adherent cells were removed, and infection was performed with *Lm*007*, Lm*N420*, Lm*TERT-/-, NAB, and TAB promastigotes in the stationary growth phase (MOI 10:1) for 3 h at 34 °C and 5% CO_2_. After 3 h, the cultures were washed with PBS, and the infection was evaluated after 24, 48, and 72 h. The infection was evaluated by determining the percentage of infected macrophages and the number of intracellular amastigotes after counting 300 Panotic-stained (Laborclin) macrophages. The percentage of infection was determined by the equation 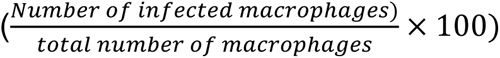. The infection index was determined by multiplying the percentage of infected macrophages by the mean of intracellular amastigotes per infected macrophage [114].

### Indirect immunofluorescence assay of *Leishmania major* infection

Samples from infected mice were tested for antibodies against *Leishmania* spp. Using the indirect immunofluorescent assay test (IIFA) [115]. The antigen for sensitization of the slides was produced using *L. major*-like antigen promastigotes (international code MHOM/BR/1976/JOF provided by Osvaldo Cruz Institute, FIOCRUZ, Rio de Janeiro, and maintained at the Zoonosis Diagnostic Service at FMVZ, UNESP, Botucatu, São Paulo, Brazil) kept in tubes containing 9 mL of liver infusion broth and tryptose medium and 5 mL of Novy-McNeal-Nicolle medium. After drying, the slides were kept at −20 °C until use. The antibodies by *Leishmania* were studied using serial serum dilutions of 1:40, 1:80, 1:160, 1:320, and 1:640. The dilution process was performed in microplates, using 190 μL of phosphate-buffered saline (PBS) (pH 7.2). Serial dilution started with 10 μl of the serum sample in the first well (1:20). Subsequently, 100 μl of PBS was added into the next five wells by pipetting a 100 μL volume from the prior dilution into the next well (after homogenization) until reaching the fifth well. The last 100 μl volume was discarded. This procedure was repeated with positive and negative control serum samples. Each serum dilution (10 μL) was pipetted into the slide wells, including the positive and negative controls. The slides were incubated inside a moist chamber at 37 °C for 30 min. Afterward, they were washed twice with PBS for 10 min and dried in the incubator at 37 °C. Anti-immunoglobulin G (IgG) specific to mice, conjugated with the fluorescein isothiocyanate, was diluted according to the manufacturer’s instructions (Bethyl Labs. / Fortis Life Sciences (USA), using Evans Blue solution with PBS (1:5) and adding 10 μl to each slide dilution. The slides were again incubated for 30 min at 37 °C in a moist chamber, washed with PBS, dried in the incubator, assembled with buffered glycerol (pH 8.5), and covered with coverslips. The slides were examined using an immunofluorescence microscope (Magnification of 40×). After reading the controls, the highest dilution of the serum for which complete fluorescence occurred of the promastigotes was considered the cut-off point, equal or superior to dilution 40 [116].

### Statistical Analysis

Data collected, unless otherwise stated, were from at least two independent experiments with at least 3 technical replicates in each assay and subjected to statistical treatment using GraphPad Prism V.9. Statistical analysis was done with either Welch t-test where two groups were being considered, two-way ANOVA or ordinary one-way ANOVA with further Dunnett or Tukey post-hoc analysis for multi-group comparisons. All graph is represented as mean ±SD, and significance was tested at P≤0.05 at all times (*p≤0.05, **p≤0.01, ***p≤0.001, ****p≤0.0001).

## Ethics statement

Animal procedures were approved by the Ethics Committee for Animal Experimentation of the Instituto de Biologia, Universidade Estadual de Campinas (UNICAMP) (protocol: 5719–1/2021) for experiments using mice.

## Acknowledgments

We thank Professors Paolo Di Mascio and Graziella E. Ronsein for the Mass spectrometry analyses performed at the Redox Proteomics Core of the Mass Spectrometry Resource at Chemistry Institute, University of Sao Paulo. We thank Jose Buratini Jnr and Thaisy Tino Dellaqua for their assistance performing the RT-qPCRs. We also thank Robson Francisco Carvalho, Florencia M. Barbé-Tuana, Theresa Teixeira, and Nuhu Osman Attah for their critical reading and suggestions regarding the project and this manuscript. We extend our appreciation to members of the Telomeres Laboratory for their diverse contributions to the project.

## Author contributions

**Conceptualization**: M.E.S. and M.I.N.C.; **Methodology:** M.E.S., B.C.D.O., H.B., D.A.S., L.C.A., R.M.B., M.M.B., M.N.C.S., B.D.M., H.L., J.I.A., A.C.C., and M.I.N.C.; **Validation:** M.E.S.; B.C.D.O.; **Formal Analysis:** M.E.S., D.A.S., R.M.B., M.M.B., M.N.C.S., B.D.M.; **Investigation:** M.E.S., B.C.D.O., H.B., D.A.S., L.C.A., R.M.B., M.M.B., B.D.M., J.I.A.; **Resources:** M.E.S., D.A.S., M.N.C.S., B.D.M., H.L., A.C.C., and M.I.N.C.; **Data Curation:** M.E.S. and D.A.S.; **Writing – Original Draft:** M.E.S.; **Writing – Review & Editing:** M.E.S., B.C.D.O., H.B., D.A.S., L.C.A., M.N.C.S., B.D.M., J.I.A., A.C.C., and M.I.N.C.; **Visualization:** M.E.S. and M.I.N.C.; **Project Administration:** M.I.N.C.; **Supervision:** M.I.N.C.; **Funding Acquisition:** M.I.N.C.

