## Supplemental file for "Ablation of telomerase reverse transcriptase in *Leishmania major* results in a senescent-like phenotype and loss of infectivity"

### **Supplementary information**

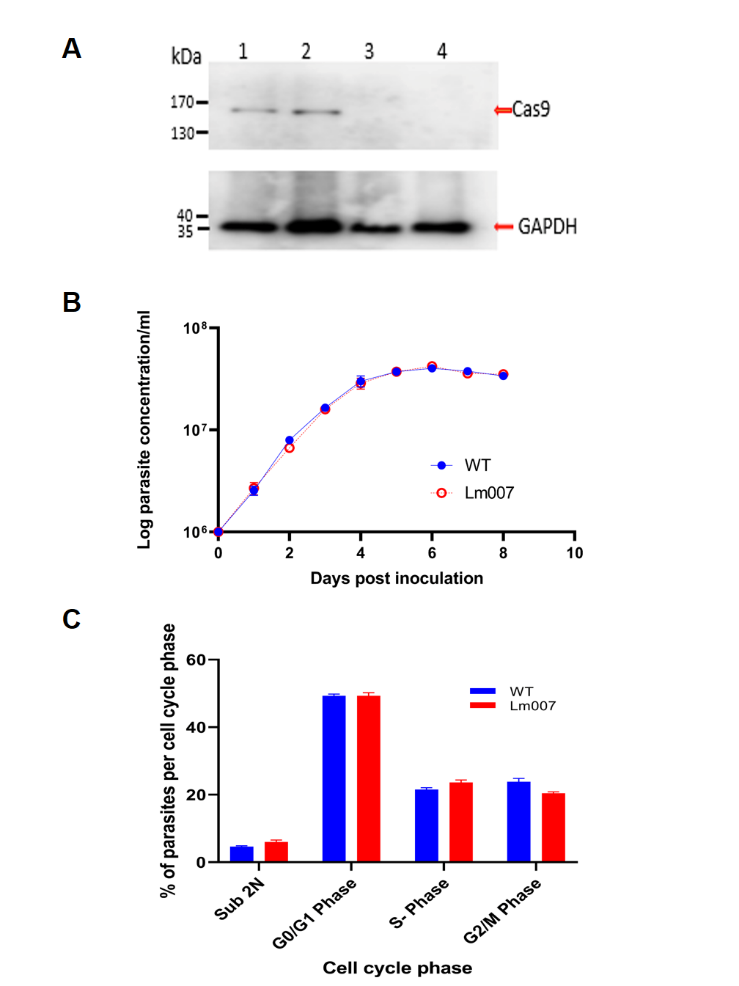

**S1 Fig. Expression of pTB007 in L. major (Lm007) does not affect the growth and cell cycle of the parasite.** (A) Western blot image for Cas9 expression validation. Lanes 1 and 2: selected Lm007 clone; Lanes 3 and 4 WT L. major electroporated and nonelectroporated, respectively. (B) Growth profile of WT and Lm007 (C) DNA content assessment of WT and Lm007.

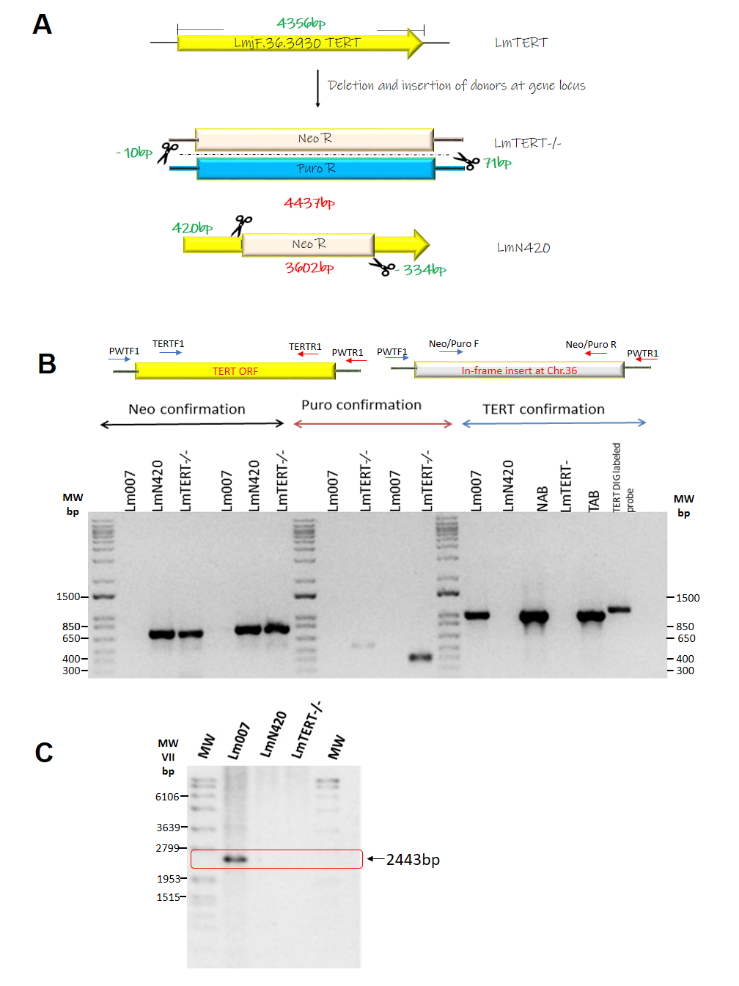

**S2 Fig. Strategies for the deletion of LmTERT and the methods of confirmation.** (A) Schematic showing the two strategies used in ablating LmTERT. The top schematic shows the full TERT gene and its corresponding size in green. The middle schematic shows the deletion of the full TERT gene (LmTERT-/-) being replaced by Puromycin (PuroR) and Neomycin (NeoR) resistance genes. The bottom schematic shows the deletion of an internal region of the TERT gene leaving 420bp upstream and 334bp downstream. The LmN420 cut site is replaced by a NeoR. The size of deleted regions in both lineages is given in red, and the cut site of the Cas 9 endonuclease relative to the start and stop codons is written in green (Middle and bottom schematic). (B) PCR amplifications confirm the successful ablation of the LmTERT and the proper integration of donor templates. Schematic maps show the positioning of primers used in the confirmation (C) Southern blot confirmation of LmTERT absence in both knockout lineages using dig labeled probe and digesting genomic DNA with XhoI.

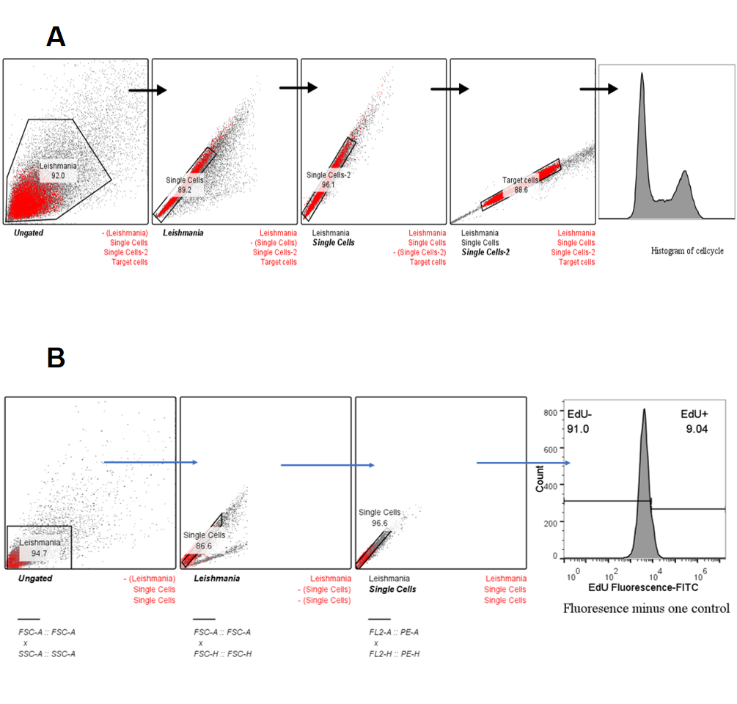

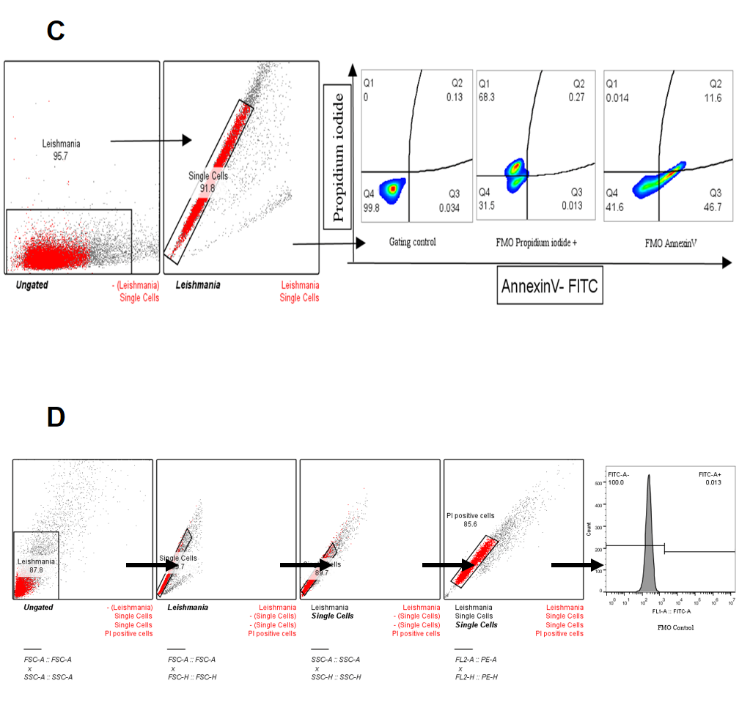

**S3 Fig. Representative gating paths used in analyzing flow cytometry data.** (A) Gating strategies used to select cells for DNA content analysis (B) EdU gating steps to select parasites undergoing synthesis and parasites that are not (C) Annexin assay gating strategies to select for cells that are positive, negative, both or neither for the stains used (Annexin V and Propidium iodide). (D) Gating paths are used to determine the percentage of DNA fragmentation. All data were analyzed using Flowjo V.10.0.7r2. The final number of cells considered for statistical treatment was within the same range for all groups considered. Fluorescence Minus One (FMO) controls were used to gate out flow-through fluorescence signals.

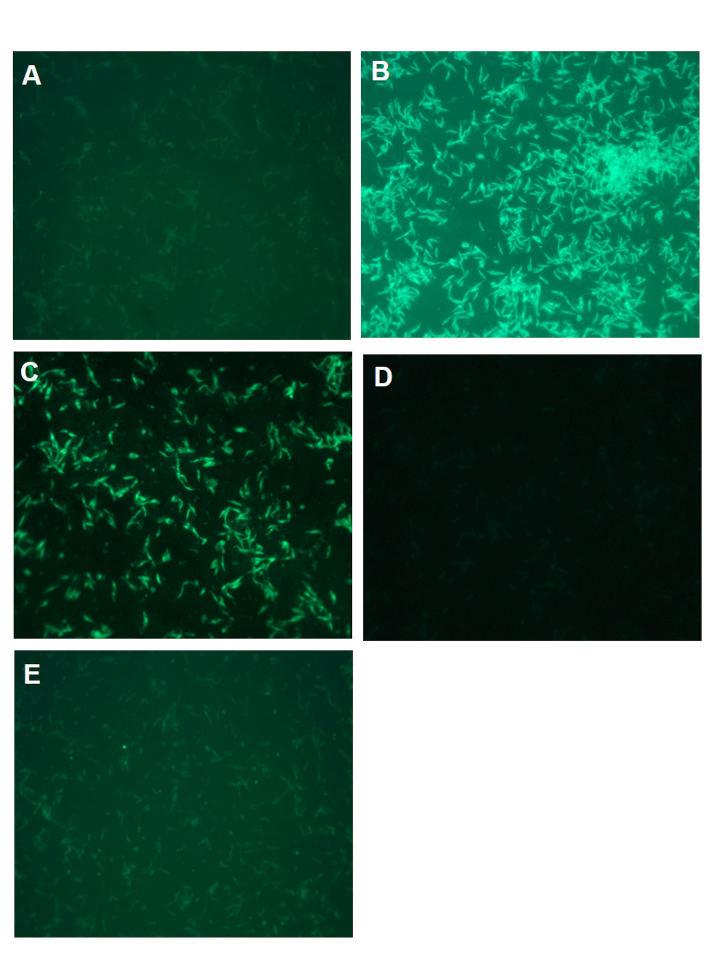

**S4 Fig. Fluorescent antibody test of Leishmania major infection in experimentally infected mice.** Images from the indirect immunofluorescence of sera were obtained from infected mice in this study. **(A)** Negative control and **(B)** positive control of the technical procedure **(C)** Results from Lm007. **(D)** and **(E)** LmN420 and LmTERT-/-, respectively.

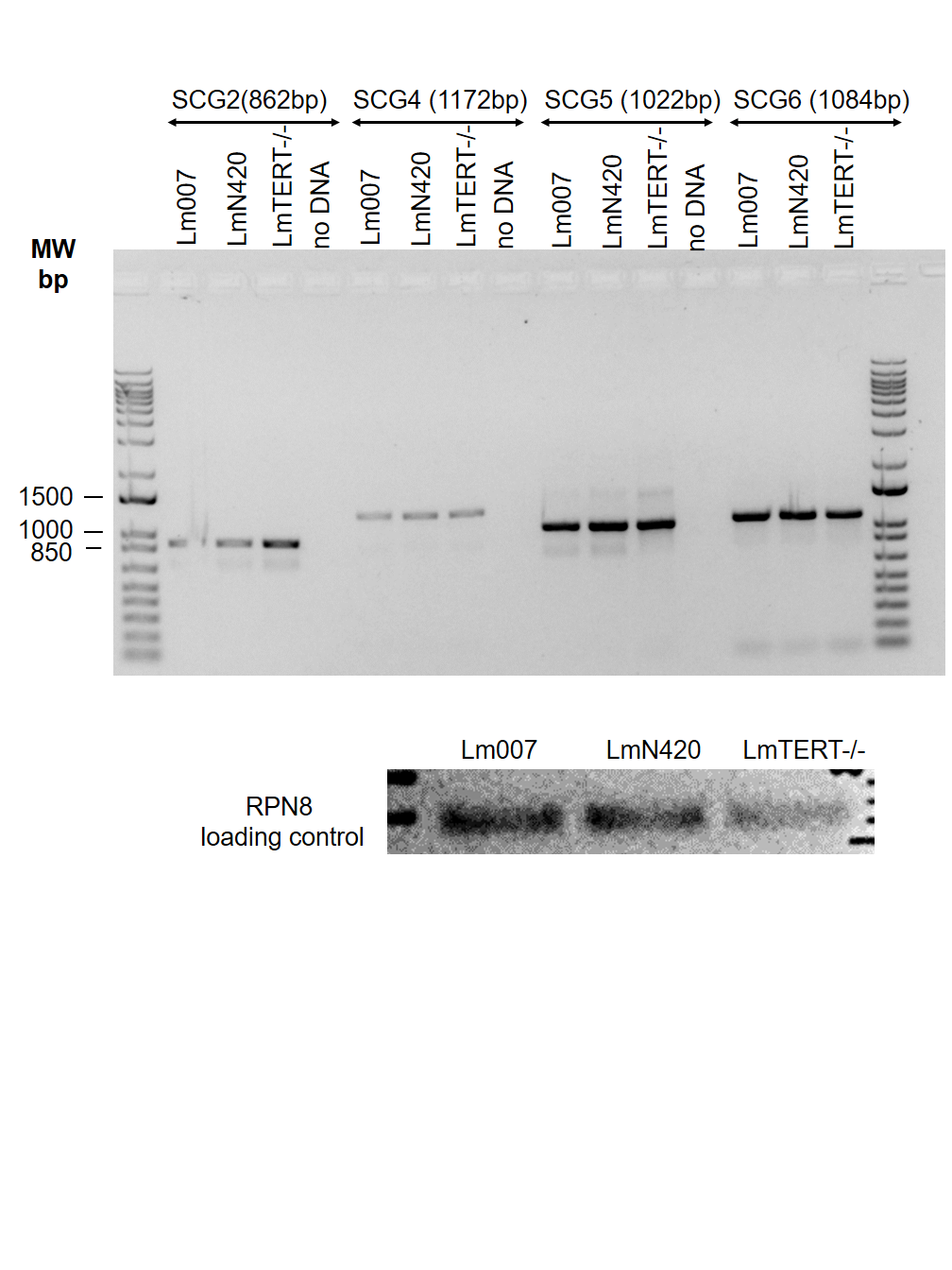

**S5 Fig. SCG gene amplifications.** Agarose gel images of the amplicons for SCG2, SCG4, SCG5 and SCG6

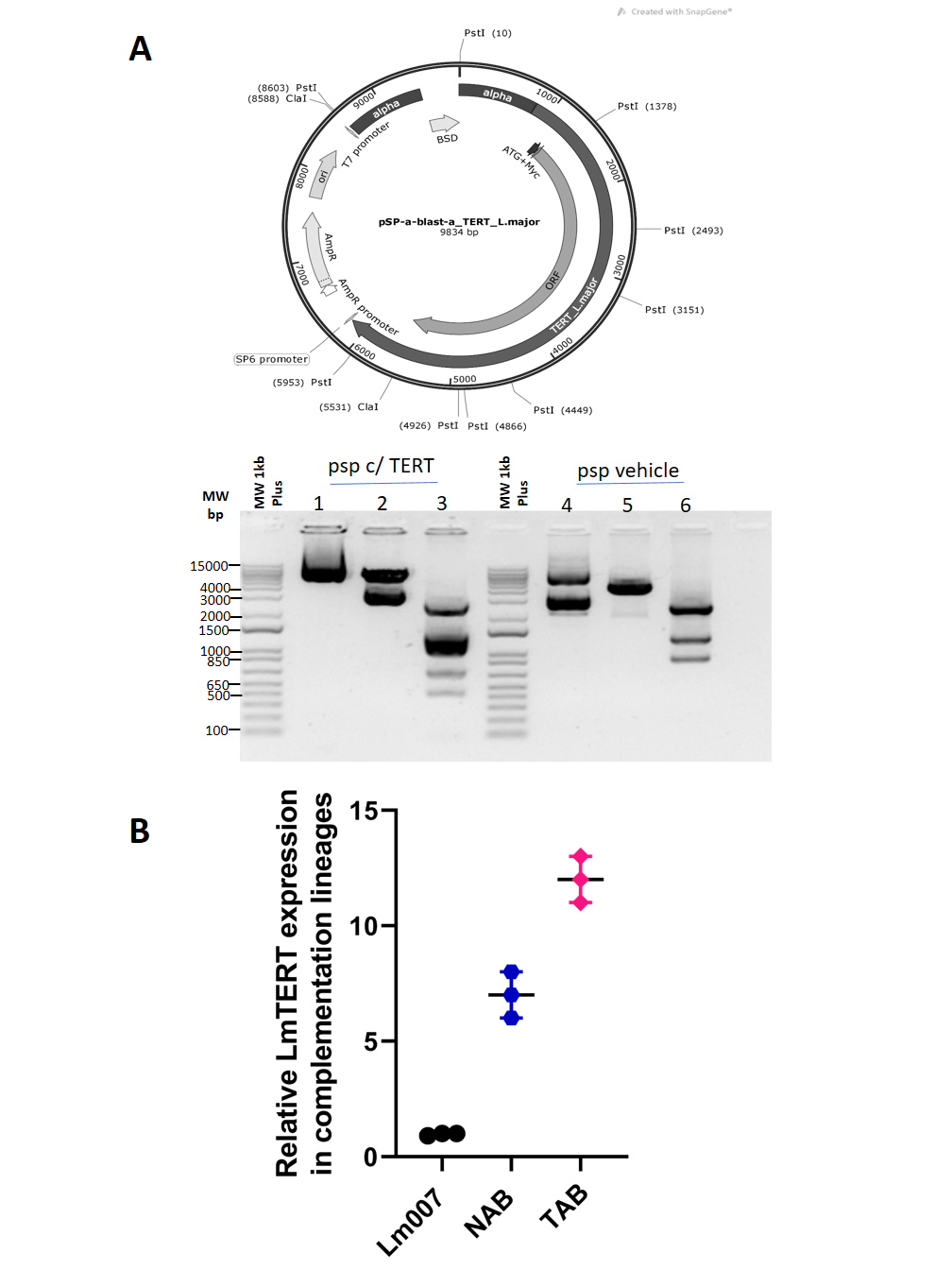

**S6 Fig. Plasmid engineering for LmTERT reintroduction in knockout lineages (complementation).** (A) Map of Psp plasmid inserted with WT LmTERT gene sequence. The map also shows the restriction sites used in restriction fragment length polymorphism confirmation. Restriction digestion of plasmid DNA to confirm the presence (left panel) and absence (right panel) of LmTERT. 1 and 4 are undigested plasmids, 2 and 5 are plasmids digested with ClaI, and 3 and 6 are plasmids digested with PstI. (B) Expression of LmTERT after complementation in knockout lines compared to Lm007.

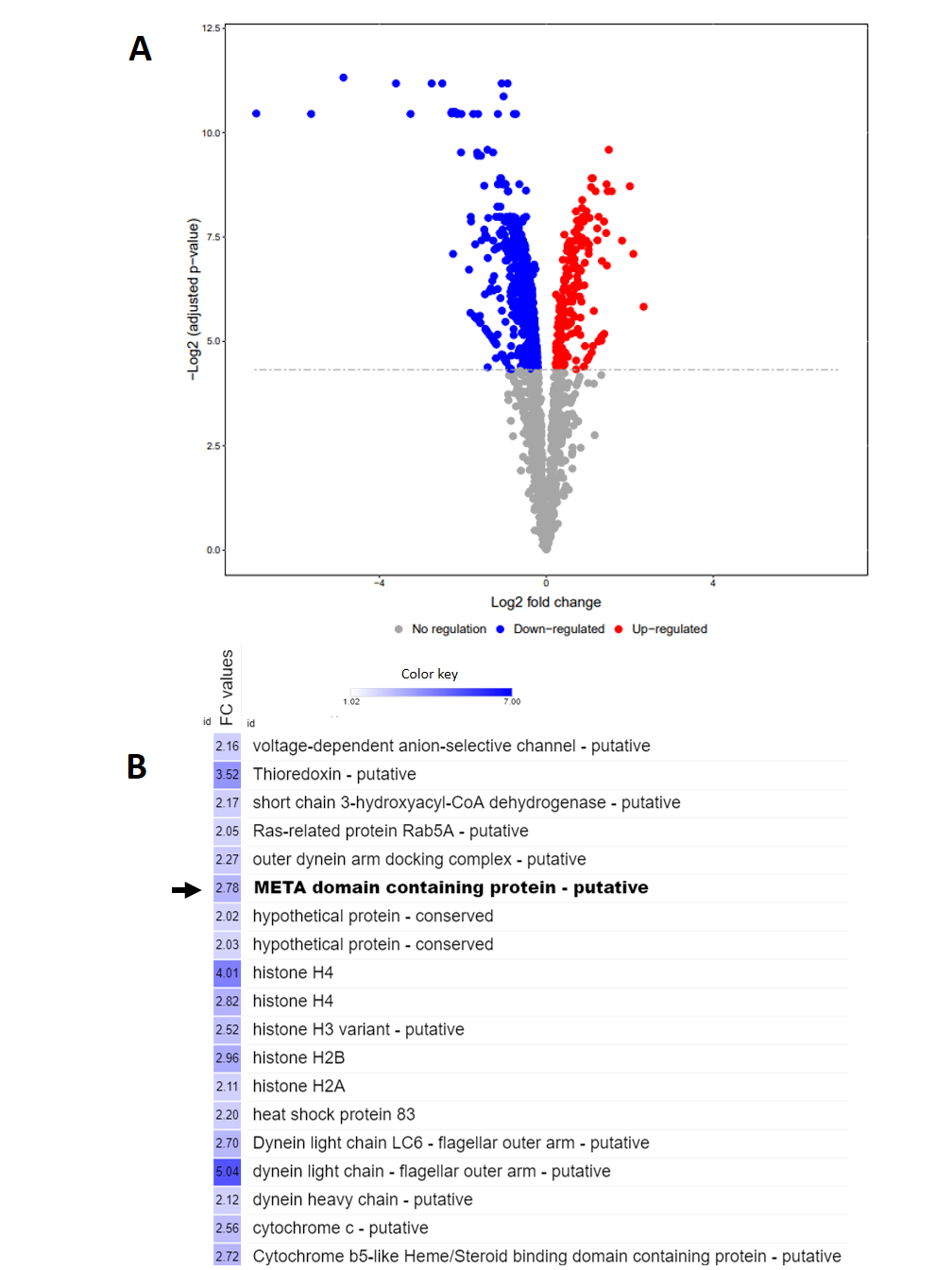

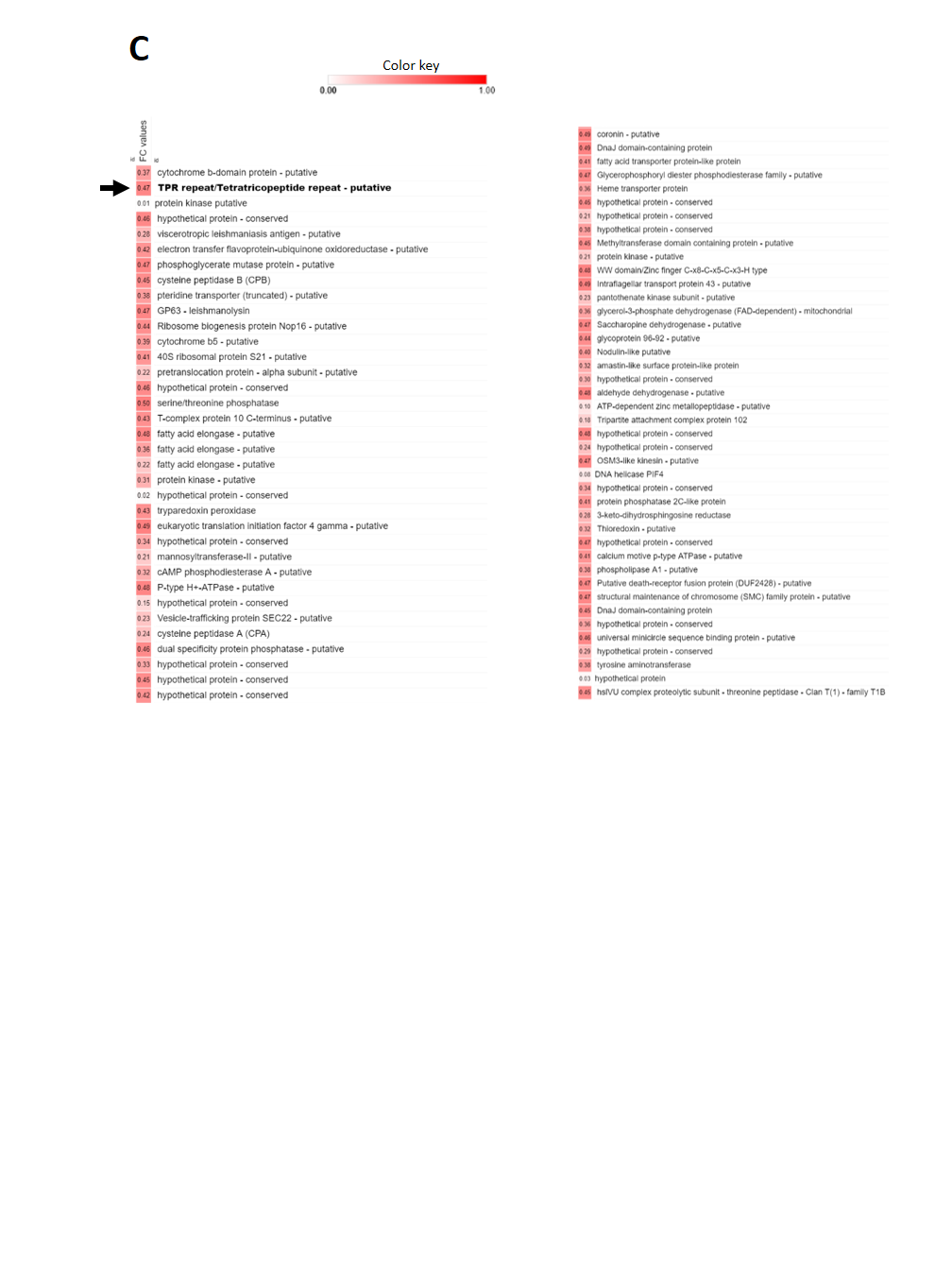

**S7 Fig. Differential protein expression. (A)** The volcano plot shows differential protein expression between Lm007 and LmTERT. **(B)** A morpheus derived heatmap depicting significantly downregulated proteins and of interest is the META domain containing protein boldly highlighted. **(C)** Heatmap of significantly upregulated proteins after TERT ablation, in bold is the TPR overexpression.

**S1 Table. List of oligonucleotides used in this study**

| **Primer Tag** | **Source** | **Identifier** |
| --- | --- | --- |
| **Oligonucleotides used in knockout procedure** | | |
| G00 (is a primer sequence, common sgRNA primer for amplification) | [39] | N/A |
| sgRNA_TERT_420 (for generating *Lm*N420):  GGAGGAGTTCTGTCGGTCGG | This paper | N/A |
| sgRNA_TERT_4022 (for generating *Lm*N420):  GGACCTCTTTAAAAGGGCGG | This paper | N/A |
| sgRNA_TERT_5'-UTR (for generating *Lm*TERT-/-):  GTACCATGAACGAGGCAAGG | This paper | N/A |
| sgRNA_TERT_3'-UTR (for generating *Lm*TERT-/-):  TAACCCCAACACTCACAGAG | This paper | N/A |
| Donor Primer (Forward):  TCCCCCCTACCAGTACCCCTCGCGTCTCCGgtataatgcagacctgctgc | This paper | N/A |
| Donor Primer (Reverse):  ACGCGGTAGATACCGAGGCACCGTCAGAGGccaatttgagagacctgtgc | This paper | N/A |
| **Oligonucleotides used in knockout confirmation** | | |
| PWTF1-(TERT 5' UTR):  GGTACGTCATAGGCACTTGAAAG | This paper | N/A |
| PWNeoF2 (NeoR ORF):  CAGACAATCGGCTGCTCTGA | This paper | N/A |
| PWNeoR1 (NeoR ORF):  AGGTCGGTCTTGACAAAAAGAACC | This paper | N/A |
| PWNeoF3 (NeoR ORF):  TGACGAGTTCTTCAGCTCCG | This paper | N/A |
| PWNeoR2 (NeoR ORF):  GAAACATCGCACACGGATGG | This paper | N/A |
| PWTR7 (TERT 3' UTR):  TGTGTGTCTGCAATCAGCC | This paper | N/A |
| Puro1 UTR 5' Rv (Puro ORF):  TTCGTGAGAGAAACCTGTAG | This paper | N/A |
| Puro2 UTR 3' Fw (Puro ORF):  AATACAGGCACGGTCCT | This paper | N/A |
| **Primers used in complementation (addbaack) confirmations (RT-qPCR)** | | |
| *Lmj*TERT2R for Target gene:  CACCCGCTCTTGTGGTAAGT | This paper | N/A |
| TERTF2 for Target gene:  CAGTTTTTGCGAGGAGGTGC | This paper | N/A |
| RPN8 F for Reference gene:  ATGAACCGCCGCAAGCT | [117] | N/A |
| RPN8 R for Reference gene:  GGCGCGCGACGACGATCTTTGATT | [117] | N/A |
| **Primers used for *SCG* genes** | | |
| SCGU3F (Forward primer used in combination with all other reverse primers-universal):  CTCGACCCTCTTCAGCTGC | This paper | N/A |
| SCG7R1 (Primer for SCG 7 confirmation):  CGCTGTGTAGTCAGAATCC | This paper | N/A |
| SCG6R1 (Primer for SCG 6 confirmation):  TTGCTGCGGCGTACTCGA | This paper | N/A |
| SCG54R1 (Primer for SCG5 and Forward primer for 4):  GAGTCCTCAGAATCATGCTCC | This paper | N/A |
| SCG4F (Primer for SCG4):  ATGCAGCCAAGCGTGTGC | This paper | N/A |
| SCG2R1 (Primer for SCG2):  CCAACACGATCAAGTGTCT | This paper | N/A |
| SCG3R1 (Primer for SCG 2):  TCGCCAAGGATGACCGCAA | This paper | N/A |
